# Multiscale active transport driven by gravitactic bioconvection promotes resilience in algal blooms

**DOI:** 10.64898/2026.06.21.733611

**Authors:** Soumitree Mishra, Jayabrata Dhar, Anupam Sengupta

## Abstract

Algal blooms are frequently dominated by motile species^1,2^ whose vertical migration enhances resource acquisition and bloom development^3,4^. Yet bloom conditions present a paradox: high cell densities intensify nutrient depletion^5^ and self-shading^6^, making individual swimming increasingly costly under severe resource limitation. How motile blooms persist and remain resilient under such stress remains unresolved^7^, particularly as climate-driven warming strengthens stratification and resource scarcity^8,9^. Here we show that the red-tide-forming phytoplankton *Heterosigma akashiwo* overcomes bloom-induced constraints through bioconvection, a self-generated active flow that emerges above a critical cell density (>1.5×10^5^ cells/ml). Using a custom *ocean-on-chip* platform that recapitulates bloom-relevant constraints, we identify an optimal synergy of cell concentration, swimming speed and gravitactic stability that promotes the formation of persistent bioconvective plumes. At constant cell density, plume onset is governed by two phenotypic traits— vertical swimming velocity and reorientation time—demonstrating that collective transport is governed by the biophysical traits of single cells. We show that bioconvection drives ecologically relevant multiscale transport, enhancing exchange of molecules and micro-cargo across stratified interfaces, mimicking transport of nutrients, extracellular vesicles^10^ and co-existing species in a bloom environment^11^. By enabling cells to hitch a hike on self-generated flows when active propulsion becomes energetically prohibitive, bioconvection-mediated transport improves nutrient delivery, restores photosynthetic performance, reverses lipid accumulation associated with nutrient-stress, and facilitates recovery of cellular motility to ultimately mitigate resource limitations. Our findings identify bioconvection as a population-level adaptive mechanism that sustains algal blooms, and reveal a previously unrecognised role of collective microbial motion in bloom persistence under ecological stresses.

**One sentence summary:** Self-organised bioconvection drives multiscale transport and resilience in algal blooms.

## Introduction

Bioconvection is a self-generated collective phenomenon^12^ in active matter systems, whereby motile microorganisms spontaneously generate patterns and large-scale flows. The onset of these active flows triggered via confluence of biological and environmental interactions^13–15^, as the microbial suspension surpasses a critical onset parameter, known as bioconvective Rayleigh number, *Ra*^16,17^. This non-dimensional parameter captures the relative effects of cell accumulation under gravity (that drives fluid motion) to the opposing effects arising due to viscous dissipation and cell diffusivity. Local accumulation of motile, stimuli-responsive cells^18–22^ in the upper liquid layers – typically 2 to 5 times larger than the mean concentration^16^ (∼ 10^5^ cells/ml), create local density differences which are sufficient to trigger hydrodynamic instability, transforming the cell-rich layers into sinking plumes and overturning them as convective rolls visualised as double-gyro structures^23^. Sustained by negative gravitaxis (swimming against the gravity vector) at the level of individual cells, this interplay of gravity and collective hydrodynamic forces drives dynamic pattern formation, ultimately emerging into stationary bioconvective rolls^14,24,25^.

In case of gravitactic bioconvection, density instability is created by negatively gravitactic cells (e.g., due to bottom-heavy or asymmetric microbes^26^), which are heavier than the surrounding fluid medium^12^, migrate opposite to the direction of the gravitational field. Such self-organised dynamic structures have been long studied^27^, across diverse groups of microbes^28–31^, revealing how directional swimming leads to emergent and sustained fluid motions which scale two to four orders of magnitude compared to the size of the individual cells driving these flows^32^. Recent studies have demonstrated the impact of these flows in natural aquatic ecosystems in nature^14,33,34^, with bioconvective mixing occurring up to meters in length scales, signifying their ecological and biogeochemical functions^34,35^. The cell concentrations required for bioconvection are similar to those observed during harmful algal bloom (HAB) outbreaks^36^ – when algal population rapidly expands, directly linking this hydrodynamic instability to ecologically relevant conditions.

Algal blooms, increasingly intensified by climate change^37,38^, pose extreme physicochemical environments to the algal population in the surface layers^7^, via increase in competition for light and nutrients. Water-column stratification, weak vertical mixing, and anthropogenic nutrient loading are widely recognized as key habitat-level factors which not only promote algal blooms, but also confer resilience to them^38–40^. The dense microbial layers – a key feature of algal blooms – can alter the physical structure and biodiversity of the upper ocean due to density stratification as well as abrupt attenuation of light intensity below the algal layers^5,6,40,41^. The density stratification offers conditions to trigger a Rayleigh–Bénard (RB) type hydrodynamic instability^42,43^, driven actively by the motile microorganisms, yet, its impact on microbial behavior and physiology, and ultimately, its role in sustaining algal blooms has been grossly overlooked. In contrast to passive RB instability where the fluid motion is sustained by an external energy source, active RB rolls are maintained by the spontaneous energy input due to the gravitactic swimmers.

Aggregation of algal cells under blooming conditions can prevent sunlight penetration to deeper layers, thereby reducing the photosynthetic activity and growth of phytoplankton lower in the water column. In aquatic ecosystems, the emergence and persistence of harmful algal blooms (HABs) are shaped by biotic and abiotic interactions^44^ and feedbacks, including nutrient enrichment. While diel vertical migration^3,45,46^ (DVM) is considered as a key biophysical trait that facilitates bloom formation, vertical transport may be constrained by the density stratification^47^, restricting both cell movement^48,49^ as well as the nutrient fluxes. Under such nutrient-limited and stratified conditions, sustained active migration may therefore become energetically prohibitive^41^.

Despite these ecophysiological constraints, algal blooms are often persistent^4,48^, and even resilient for extended periods under resource-limited settings. For bloom-like concentrations (∼10^3^ 10^6^ cells/ml)^48,50^ of gravitactic cells, could then self-generated, collective dynamics – a biophysical factor that remains unexplored – contribute to enhancing nutrient and light availability, and thereby confer biophysical resilience to an otherwise resource-limited bloom?

Under ecologically relevant cell concentrations of a bloom forming species, we demonstrate that, reduction in the nutrient concentration triggers an interplay between optimal cell concentration (> 10^5^ cells/ml) and gravitactic behavior, that ultimately promotes the onset of self-generated bioconvective flows. Under nutrient replete conditions, bioconvection was absent. Our results reveal that bioconvection occurs within an optimal range of swimming speed and cell reorientation timescales, supporting cells to hitch a ride on such active flows, as the cell-level propulsion becomes energetically expensive due to nutrient- and light-limited conditions. Transport of micro-cargo, spanning 2 μm to 20 μm in size, served as a proxy for diverse particulates observed in a bloom, including non-motile diatoms, weakly motile, or stressed flagellates, as well as extracellular vesicles (EVs) which mediate intercellular signaling^10,51^, biogeochemical exchange^52^ and bloom dynamics^53^. Furthermore, stationary bioconvective flows sustain enhanced molecular transport across the density barrier, highlighting their significance in natural ecosystems. Finally, by quantifying the photosynthetic performance, and the accumulation of intracellular lipids, we confirm that bioconvection-mediated enhanced transport mitigates nutrient deficits and recovers photosynthesis, thereby improving the cellular physiology, alongside aiding phenotypically weaker swimmers back to their motile state. Taken together, bioconvection confers an adaptive resilience under blooming conditions, underscoring an important, yet so-far-overlooked, emergent fluid motion that offsets ecophysiological constraints and sustains algal blooms.

## Results

### Phenotypic traits govern the onset and dynamics of gravitactic bioconvection

Using *Heterosigma akashiwo* (CCMP3374, hereafter HA3374), a red tide forming species, we show that an interplay of cell concentration (depending on the physiological growth phase) and phenotypic traits, govern the onset and dynamics of self-organised bioconvection (Fig. 1a, Supplementary Movie 1). This collective-scale convection arises when cell accumulation in the upper layer becomes unstable due to density stratification, leading to the formation of descending plumes. We observe steady bioconvective patterns, starting from the early stationary growth phase (> 200 h after fresh innoculation^26^ ), and persisting until the onset of the death phase (Fig. 1b, S1 a). The maximum cell concentration, ∼1.8×10^5^ cells/ml, is similar to dense accumulations in natural algal blooms. Appearance of these large-scale dynamics under the nutrient-limited settings enable cell populations to migrate distances of orders of magnitude larger than their individual cell dimensions, thus facilitating energetic conservation, which otherwise would have been expended due to self-propulsion. Along the physiological timescales, we capture the cell-level geometry, together with the dynamics of bioconvection via particle image velocimetry (PIV, Fig. 1c, S1 b). We found that the onset of these active flows is more likely to occur at the late stationary phase, and was absent during the late exponential stages of growth (inset-Fig. 1b (bottom panel), Supplementary Movie 2), despite both growth stages having the similar mean cell concentrations.

**Fig. 1:**
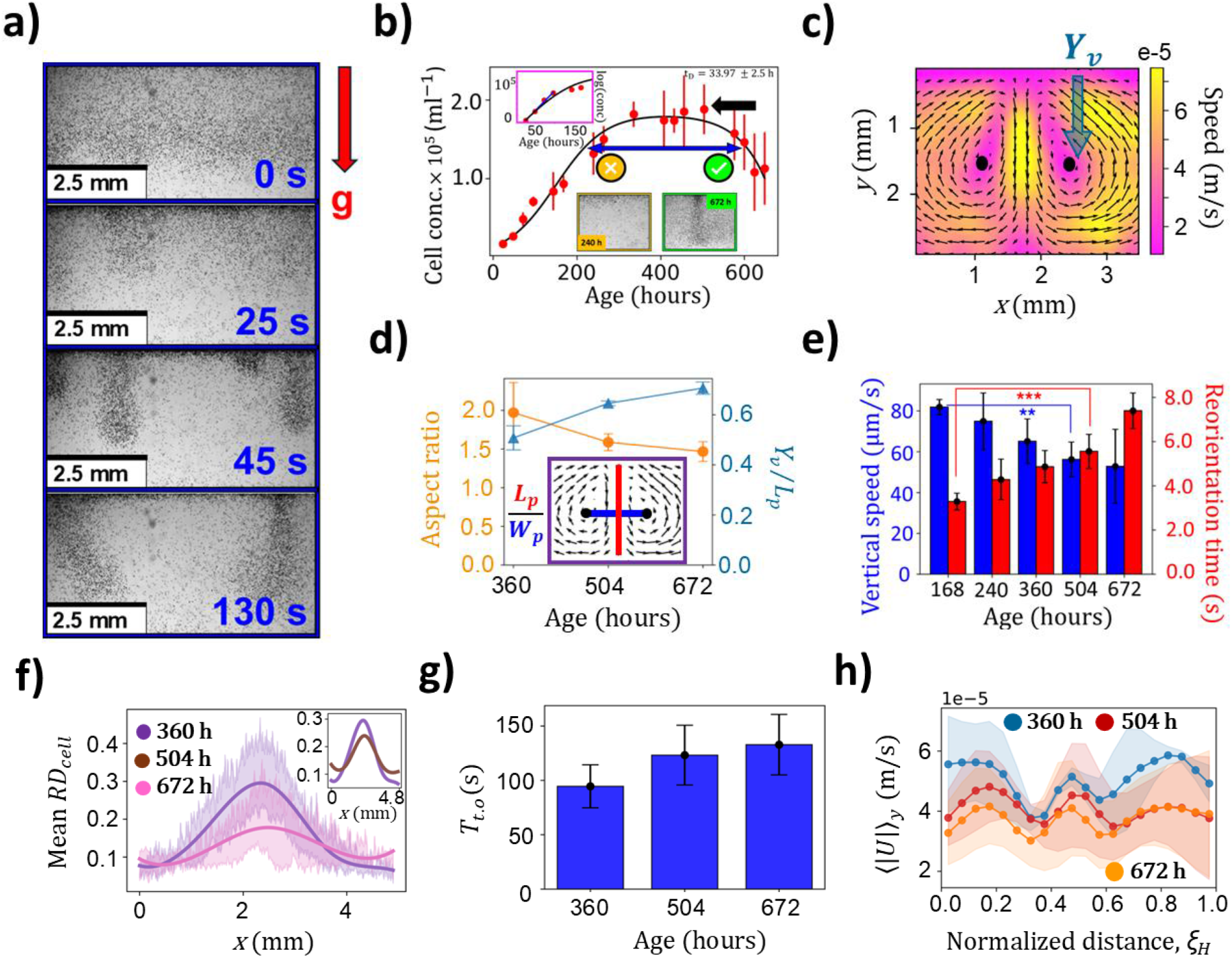
Phenotypic trade-offs regulate the dynamic plume geometry. **a)** Temporal evolution of bioconvective patterns, showing the evolution of the plume morphology over experimental timescales (Supplementary Movie 1). **b)** The growth curve of HA3374 (Black lines represent logistic fit), with the mean cell concentration (red dots) and error bars (±1 s.d. from the mean of 4 replicates). The arrow marks the time point for experimental observation. The top inset plot shows the semi-log distribution of mean cell concentration over time. Bottom inset panel shows the differences in bioconvection with same cell concentration but at two different growth phases (cells at earlier growth stage, failed to show bioconvection – Supplementary Movie 2). **c)** Velocity field of a bioconvective plume extracted using particle image velocimetry (PIV). The vortex height (*Y*_*v*_) measures the distance between the top plane and the circulation center. **d)** Aspect ratio and the normalized vortex height, of bioconvective plumes plotted across different growth stages (3 biological replicates), revealing changes in plume structure as cells progress through their life cycle. Inset plot indicates the plume length (in red) and the plume width (in blue) for computing the aspect ratio. **e)** Vertical swimming speed and reorientation time of individual cells in quiescent fluid environment across different growth phases, reflecting their phenotypic traits. Bar plots show mean values with standard deviations from 6 replicates. Asterisks denote statistically significant differences (p < 0.01). **f)** Mean relative cell density of a stable bioconvective plume across different growth phase. Broader distributions indicate diffused plumes arising from cellular ageing, higher diffusivity, and reduced activity. Shaded regions represent standard deviation from three biological replicates. **g)** Bar plot illustrating the turnover time for swimming cells in bioconvection. Error bars represent ±1 standard deviation from three biological replicates. **h)** Mean velocity magnitude averaged along the vertical (*y*) direction projected along a normalized horizontal plane. Shaded regions highlighting standard deviation from three biological replicates.

We quantify the geometry of a stable plume by estimating its aspect ratio, the ratio of the plume length (*L*_*p*_) to its width (*W*_*p*_), extracted from the PIV data (Fig.1d, inset). As the population ages, the aspect ratio decreases from 1.97± 0.39 at 360 h to 1.46 ± 0.12 at 672 h (P = 0.042 < 0.05), significantly broadening the bioconvective patterns (Fig.1d, S1 b). During the same period, the vortex center (in the vertical plane) shifts downward significantly, from 0.51 ± 0.05 to 0.71 ± 0.03, the shifts normalised to the plume length (p = 0.0079 < 0.01). This suggests a delay in the collective-scale turnover of older populations (Fig .1d). Under these conditions, phenotypically weaker cells, i.e., cells with lower swimming speeds and longer reorientation times get passively advected by the self-generated bioconvective flows, thus sinking to greater depths before being turned over by the collective circulation, resulting in a downward shift of the vortex center. During the early growth phases, younger cells swim faster (81.9 ± 3.7 µm/s), relative to the cells from the later stages of growth (52.8 ± 18.1 µm/s, Fig. 1e, S1 d). In parallel, cells actively reorient back to their equilibrium swimming direction when perturbed: the corresponding timescale is a measure of the gravitactic stability^25,26^, signifying how quickly a cell returns back to its stable alignment (Fig. S2 b, Supplementary Movie 3). Consequently, during the early phases of growth, the combination of high cell motility and low reorientation timescales (Fig. S2 e) disrupts the gravitational instability by escaping from downwelling plumes, decreasing the likelihood of plume nucleation and a stable bioconvection. As the nutrient availability drops, the reorientation time goes up from 3.28 ± 0.38 s (at 168 h) to 7.39 ± 0.80 s at 672 h (Fig. 1e, S3), leading to spatially dispersed mean concentration distributions within stable plumes formed by phenotypically weaker populations (Fig. 1f, S2 e), in agreement with the geometrical analysis of plume structure. Leveraging the PIV data, we numerically compute the timescales of the net turnover by bioconvective plumes (Fig. 1g, S1 c, Supplementary Movie 4), we show that longer turnover timescales – achieved via lower gravitactic stability and reduced vertical speeds – drive the nucleation of stable bioconvective plumes during the later stages of growth. This observation was further confirmed by the depth averaged velocities, which decline on average for plumes formed by phenotypically weaker populations (Fig. 1h). During the mid-exponential phase, the optimal balance of traits creates a favorable condition for nucleation, enabling accumulating cells in top layer to sink as plumes. In bioconvective flows, vertical transport is dominated by gravitational sinking, thereby cells expend less energy in swimming actively across depths. Bioconvection-mediated advection enables cells to collectively migrate, providing a selective advantage in nutrient-limited environments and improving access to light for photosynthesis during the daytime, when bioconvection is more dominant (Fig. S1 e).

### Cellular phenotypic traits govern micro-cargo transport due to bioconvection

To quantify emerging hydrodynamics due to bioconvective plumes, we introduce passive tracers (Fig. 2a) and analyzed their dynamics using particle tracking (Fig. S4 a-d, Supplementary Movie 5). These tracers, approximately 1/10^th^ the size of an individual cell, not only provide the local flow-field characteristics in bioconvection but also serve as a proxy for ecologically relevant micro-cargo, e.g., non-motile picoplankton (< 2 µm), or phenotypically weaker or stressed cells in the local environment which may benefit from the emergent transport. In control experiment (no cells), the mean squared displacement (MSD) scaled as ∼*t*^α^, with α = 1.19, indicating a slow passive sedimentation under gravity; while the MSD in presence of bioconvective flow varied spatially, depending on the location of the particles relative to the plume center. The central region of the bioconvective plume (Region I: the space between the two vortex centers), captures the gyrotactic focusing of downwelling cells, where microscale transport is super-diffusive with MSD ∼ *t*^1.96^ (Fig. 2b, S4 e). The region II (near region) represents the upwelling section further away from the plume center, where collective upward migration of cells was dominant. The exponent, α ∼1.86, suggests presence of an advective transport component (Fig. 2b, S4 e). When larger micro-cargo (20 µm mean diameter, comparable to the cell size) was introduced, a notable reduction in the cargo transport was observed with a reduced MSD ∼ *t*^1.72^ as compared to the smaller micro-cargo in the region II (Inset Fig. 2b, S4 g). These observations suggest that passive particles comparable to the cell body length can still undergo enhanced transport, although the MSD decreases with increasing cargo size. Such emergent transport processes could be ecologically beneficial for non-motile microbes living in the community, stressed phytoplankton with reduced motility, or for transport of EVs as chemical messengers.

**Fig. 2:**
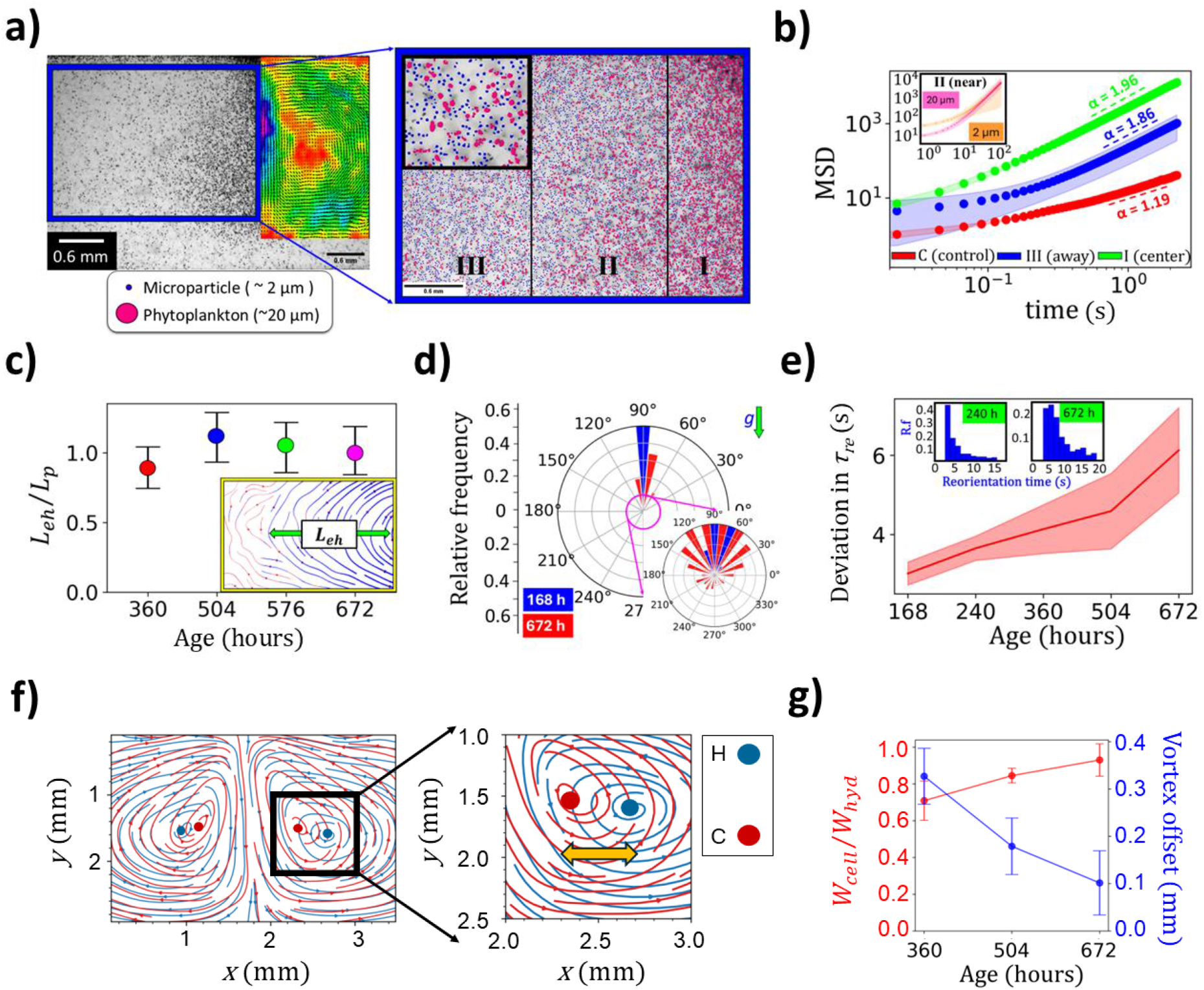
Gravitactic stability in the population regulates emerging hydrodynamics. **a)** Passive tracers (blue) are incorporated into a steady state bioconvective plume, identified through image segmentation (Supplementary Movie 5). HA3374 cells are marked in magenta. The inset provides a magnified view of segmented regions. The plume is divided into Regions I, II, and III, each spanning ∼1.2 mm in width. **b)** The slope of the mean squared displacement (MSD) versus time (*t*) shows increased super-diffusivity near the plume center (mean ± 1.s.d, 3 biological replicates). The inset plot shows the MSD for two different seized tracers (2 µm & 20 µm) analyzed in same location - region II. **c)** The length scale of enhanced hydrodynamic (*L*_*eh*_) transport, normalized by plume length (*L*_*p*_), varies with phenotypic traits at different growth stages. Error bars indicate mean ± 1 standard deviation from 4 replicates. **d)** Gravitactic direction at two different stages of growth: Magnified polar plot shows the emergence of subpopulation with random swimming direction at later stages of growth. **e)** The mean and standard deviation of noise in reorientation time within the population. Older cell cultures show a relatively larger deviation in reorientation time, *τ*_*re*_, due to higher phenotypic variability. **f)** Streamlines of actively swimming cells (red), and the emergent hydrodynamic flow (blue), were segregated and analysed separately (Supplementary Movie 6). The zoomed-in view shows that the center of the vortex of the hydrodynamic field (blue) is offset from that of the active cell circulation field (red). **g)** Active cell plume width, normalized by hydrodynamic plume width, plotted over different across generations (red). The *y*-axis (right hand) presents the vortex center offset over three biological replicates (blue).

Using PIV, we quantify the spatial range of the bioconvection-mediated enhanced transport, computed via velocity thresholding (Materials and Methods, Fig. S5 b). Figure 2c shows the range of the enhanced hydrodynamic transport, *L*_*eh*_, across the physiological growth stages. Despite the larger plume aspect ratio, younger populations (age 360 h) show reduced hydrodynamic transport as the cells escape the gyrotactic focusing due to their ballistic swimming behavior against the gravity direction (Fig. 2d, S5 c). Cells with higher gravitactic stability thus contributes relatively less to the ambient convection collectively. While the older cells (age 672 h) lose directional swimming, on average they exhibit more frequent local reorientation, consistent with greater variation in reorientation times across the population (Fig. 2e, S5 d). Thus, the gravitactic reorientation timescale becomes a key control parameter: slower the reorientation dynamics, the cells are advected more strongly into the downwelling region (before they can re-align against gravity). So, between these two limiting cases, optimal motility and gravitactic reorientation act together to strengthen gyrotactic focusing, leading to maximal hydrodynamic transport. To verify this, we delineate the streamlines arising from the emergent hydrodynamic field from those from the bioconvecting cells (Fig. 2f, Supplementary Movie 6). This reveals that the vortex offset – defined as the difference between the circulation centers of hydrodynamic flow field and bioconvecting cell field, is higher for younger populations, indicating that the collective turnover by cells happens faster than the hydrodynamic turnover. The vortex offset decreases significantly (P = 0.012) with culture age (Fig. 2g, S6 a), due to reduced active swimming against self-generated hydrodynamics, leading to closer alignment of cell motion with the passive tracer fields (Fig. 2g - blue, Supplementary Movie 6, 7). Broadening of the bioconvecting cell-plume width (*W*_*cell*_), relative to the hydrodynamic plume width (*W*_*hyd*_), from 0.71 ± 0.11 at 360 h to 0.93 ± 0.09 at 672 h, reflects an overlapping of the streamlines as cell become phenotypically weaker (Fig. 2g - red, S5 e).

### *Bloom-on-a-chip* reveals molecular trap-and-transport by the self-generated flows

The gyrotactic motion of cells creates structured flow patterns in aquatic environments, facilitating the distribution of molecular patches for population-wide benefit^29^.We quantify the spatio-temporal molecular field using a combination of numerical simulations and experimental observations wherein a chromatic dye was employed as a molecular cargo. In presence of bioconvection, the plume becomes entrained and trapped within the plume vortices, forming a localised high concentrated region (Fig. 3a, Supplementary Movie 8). Our experimental observation is confirmed by simulations, further revealing that molecular trapping and transport depends on the molecular diffusivity values (Fig. 3b). Using Finite Time Lyapunov Exponent (*FTLE*) analysis^43^, we identify entrainment zones and transport barriers (Fig. 3c, S6 b-d, Supplementary Movie 9). The mean Lyapunov exponent, *LE*, for steady bioconvection plumes (e.g., cell population corresponding to 504 h) have a narrow distribution as shown in Fig. 3d, *LE* = 0.040 ± 0.025, signifying efficient trapping. Weakly bioconvecting populations or populations lacking bioconvection exhibit broader *LE* distributions with higher means (0.048 ± 0.020 and 0.052 ± 0.024, respectively for 360 h and 642 h), indicating weaker molecular trapping.

**Fig. 3:**
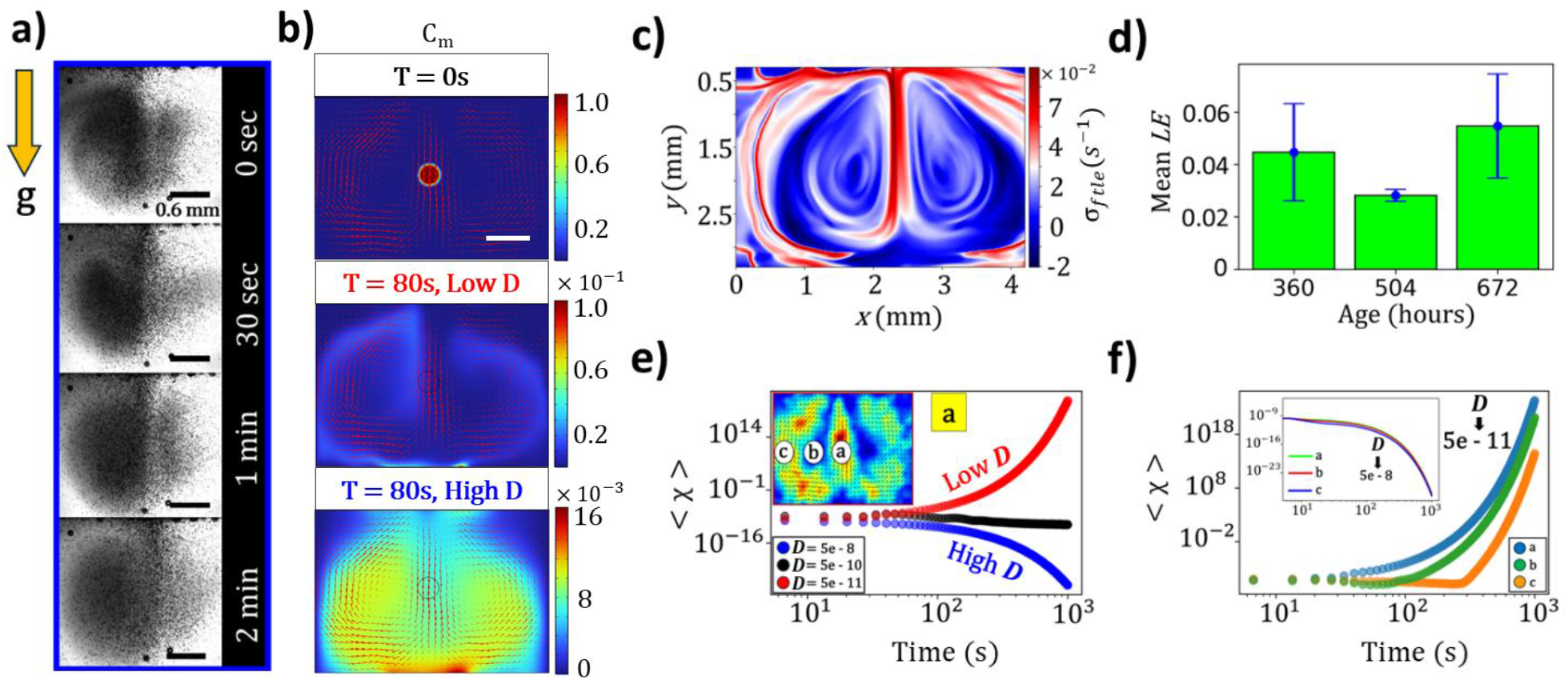
Molecular trapping and transport by bioconvection. **a)** Experimental observations over a 2-minutes window capture the trapping and transport of dye molecules (visualised in grey hue) driven by the bioconvection plume (Supplementary Movie 8). **b)** Numerical simulations, based on the experimental flow data, quantify molecular concentration over time (Supplementary Movie 10,11). Top panel: Initial location of molecular patch at the plume center (scale bar – 0.5 mm); middle panel shows the molecular distribution for a high diffusivity molecule (*D* = 5 × 10^−9^ m^2^/s); bottom panel shows bioconvective transport of a low diffusivity molecule (*D* = 5 × 10^−10^ m^2^/s). **c)** The finite time Lyapunov exponent (*FTLE*) field, computed numerically from the experiments, with the closed ridge lines, the Lagrangian coherent structures (*LCS*) delineate zones of high molecular trapping within the flow. **d)** The mean Lyapunov exponent, computed across the two-dimensional space, quantifies the strength of molecular trapping for bioconvection driven by cells across growth stages. **e)** Time-averaged scalar dissipation rate of a molecular patch positioned at the plume center, highlighting the influence of diffusivity on molecular transport dynamics (initial release location fixed at “a”). The inset presents three different initial release locations of the molecular patch within the plume region. **f)** The dependence of the scalar dissipation rate on the initial release location of a patch of low diffusivity molecules. Inset shows the initial release location-dependent dissipation rate for a high diffusivity molecules.

We solve the convection–diffusion equation on the experimentally obtained velocity field and quantify mixing using the time-averaged scalar dissipation rate ⟨*χ*⟩ through simulation (see Methods). Both the initial position of molecules and the diffusivity *D* influence the molecular transport dynamics (Supplementary Movies 10 and 11). For a single plume, we consider three initial release locations for molecules (Fig. 3e, inset) and three representative values of diffusivity, corresponding to small molecules (high *D*) and larger molecules (low *D*). For high *D* molecules, ⟨*χ*⟩ is independent of initial release locations, decaying monotonically in time (Fig. 3e). The high *D* molecules smear across streamlines and tend to replenish convergent zones around the downwelling jet, thereby entraining in the proximity of the plume’s core. This suggests efficient mixing and homogenization within plume boundary. In contrast, at low diffusivity, the dissipation rate ⟨*χ*⟩ either remains constant or increases with time, reflecting poor mixing. Low *D* molecules stretch into thin filaments, and get expelled from the vortex core due to the strong downwelling flux, and get redistributed into the surrounding fluid, thereby accumulating away from cell-rich plume regions (Fig. S7). In the context of nutrient molecules, the downwelling regions thus act as a nutrient sink, creating local regions of low molecular concentration while maintaining strong gradients across the plume axis and the boundary. This allows further transport of the nutrients within the plume core. However, ecologically-relevant low diffusivity molecules may get expelled from regions of strong vorticity or strain, as advection dominates and the weak diffusive transport is insufficient to keep the molecules entrained. Taken together, the location where the molecular patches arise, together with their diffusivities become a key determinant in shaping the transport dynamics of molecules in the vicinity of bioconvective flows (Fig. 3f, Supplementary Movie 11).

### Bioconvective transport in resource-limited algal blooms promote physiological and behavioural recovery

In natural aquatic environments, nutrient transport from deeper waters to the surface is often hindered by a density barrier formed by freshwater runoffs and elevated surface temperatures^54,55^. This stratification limits the upward movement of nutrients, creating nutrient-poor surface layers, once nutrients are depleted due to bloom. To mimic these conditions, we prepared a stratified water column with nutrient-rich, high-salinity bottom layers (42 PSU) and nutrient-poor, low-salinity surface layers (36 PSU) filled with cells. Experiments were conducted under two scenarios: one with a high cell concentration in the surface layer to enhance bioconvection and another with low cell density to avoid cell aggregation and, consequently, bioconvection (Fig. 4a). Using chromatic dye to visualise bioconvection-induced transport, we simultaneously track the size of intracellular lipid droplets. After 11 h, the chromatic dye mixes into the surface layer, breaking stratification entirely by 24 hours (Fig. S8 a,b). In contrast, diffusion-driven mixing – in absence of bioconvection – in the control setup was minimal, with the dye largely confined to the bottom of the chamber. This confirms that bioconvection can disrupt density stratification as well as enhance nutrient transport.

**Fig. 4:**
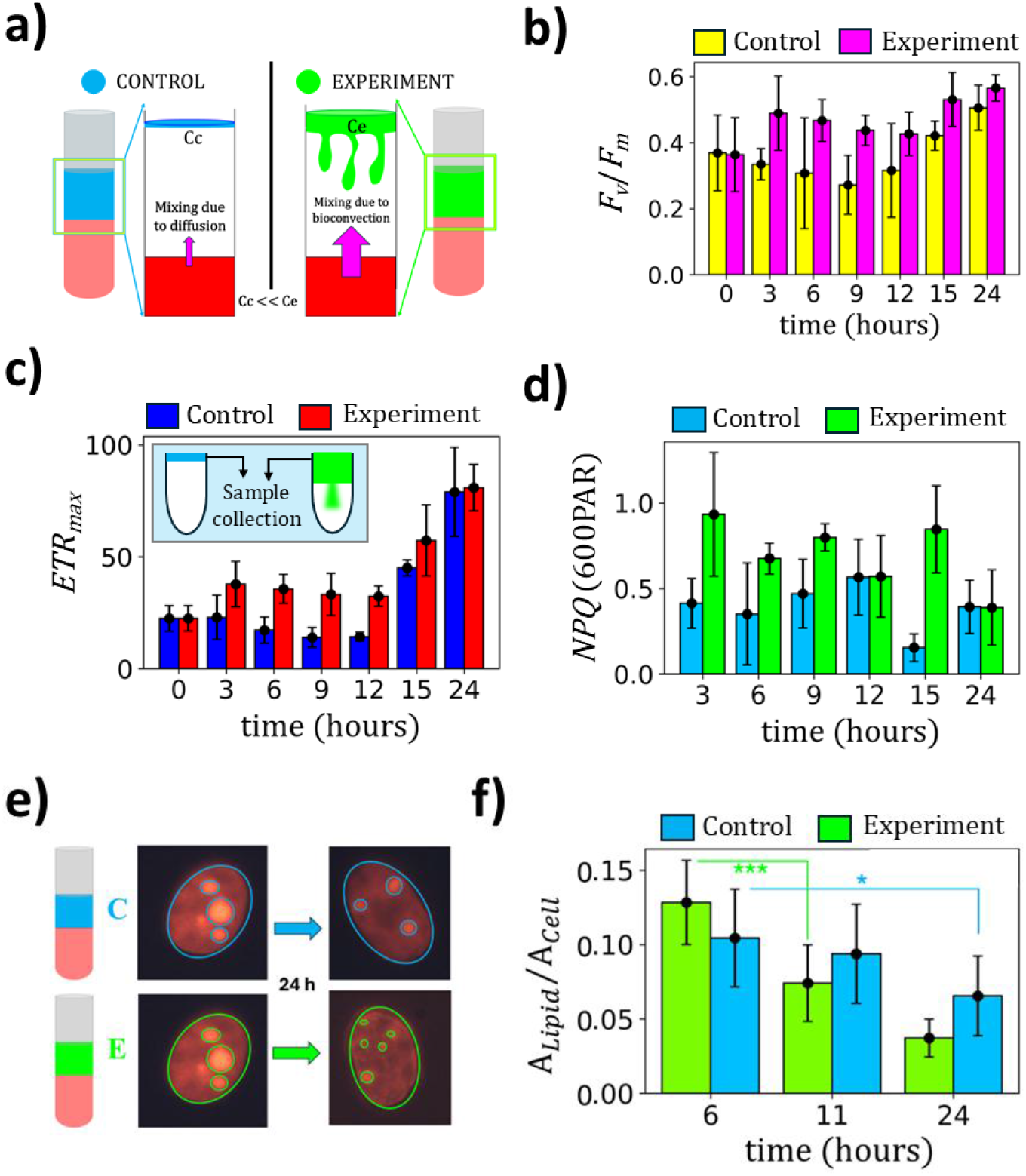
Bioconvection enhances nutrient transport and drives physiological and behavioural recovery of cells. **a)** Schematics of the control and experiments to track nutrient transport due to upwelling in stable stratification. High cell concentration at the top eventually descends as plumes, which was absent in control setup (diluted cell culture). **b, c)** Photophysiology, measured as *F*_*v*_*/F*_*m*_ and Electron transport rate (*ETR*), shows quicker recovery for populations experiencing bioconvection (inset panel Fig. c shows experiment schematics). **d)** Non-photochemical quenching (*NPQ*) for both control and experimental samples. **e)** Fluorescent signals from intracellular lipid droplets, visualised using Nile Red dye. **f)** Lipid area vs cell area drops significantly faster under bioconvection (due to enhanced nutrient distribution), compared to the control case in absence of bioconvection (transport is solely via diffusion).

To examine the physiological impact, we track lipogenesis (fluorescence imaging) and the photosynthetic performance of cells measured using PAM fluorometry^56^ (Fig. S8 c). Under nutrient limitation, lipid accumulation (Fig. S8 d-f) in cells is accompanied by a drop in the photosynthetic efficiency. However, nutrient supplementation reversed these effects, enhancing photosynthetic activity^56^. Our experiments demonstrated that cells in bioconvecting environments recover significantly more rapidly, indicating faster and efficient nutrient distribution, resulting in rapid improvement of photo-physiology (Fig. 4 b, c). The ratio of variable fluorescence (*F*_*v*_) to maximal fluorescence (*F*_*m*_) of chlorophyll (*F*_*v*_/*F*_*m*_) serves as a metric for evaluating photosynthetic efficiency and cellular physiology, measured using PAM fluorometer^42^. Within the first three hours, cells in bioconvecting samples exhibited an increase in *F*_*v*_/*F*_*m*_ (Fig. 4b) and maximal electron transport rate (*ETR*_*max*_), (Fig. 4c) indicating an accelerated nutrient upwelling from the bottom layers. In contrast, control samples, where nutrient availability relied solely on diffusion, show negligible change. Conversely, diffusion-limited cells exhibit slow recovery, due to insufficient nutrient supply. Non-photochemical chlorophyll fluorescence quenching (*NPQ*), capturing the dissipation of excess absorbed light energy as heat (∼ 600 PAR light intensity typically observed at the ocean surface), was high after 3 h, indicating an enhanced ability to mitigate light-induced stress while maintaining overall photosynthetic efficiency. Finally, cells exposed to bioconvection exhibited a rapid reduction in the lipid-to-cell area ratio, decreasing significantly from 0.13 ± 0.03 to 0.07 ± 0.02 within a span of 6 hours (Fig. 4f). This rapid lipid breakdown (Fig. 4e, f, S9) indicates efficient nutrient uptake and utilization, facilitated by bioconvection, which was not the case in control samples. Cells in a diffusion-limited environment (control case) displayed a slower lipid degradation rate, with changes in the lipid-to-cell area ratio showing less statistical significance. This suggests that nutrient supplements in these conditions is less efficient, underscoring the role of bioconvection in accelerating nutrient availability and physiological recovery (Fig. S9, S10).

## Discussion

Photosynthetic microorganisms naturally exhibit diel vertical migration (DVM) to access well-lit photic zone during the day and nutrient-rich bottom layers during the night. In natural ecosystems, they leverage this migrating behavior to shuttle between resource rich layers to form intense blooms^3^. The species we investigated in this study are highly motile under nutrient-replete conditions, i.e., during the exponential growth phases. Strong activity however suppresses the onset of bioconvection, with no visible signatures of large-scale coherent flows (Supplementary Movie 2). However, as nutrient turns limiting with increasing cell concentrations, HA3374 reached bloom-like conditions, showing weaker phenotypic traits (both in terms of vertical swimming speed and reorientation times). Because under changing nutrient landscapes, the behavioral and physiological changes serve as adaptive strategies underpinning energy cost optimization as well as survival under stressful conditions, vertical migration driven by active flagellar propulsion becomes energetically unfavorable. We show that, at later growth stages when cells are increasingly, a combination of slower motility and optimal reorientation timescales drives the onset of bioconvection. This is because, reduced motility increases the residence time of cells within unstable dense patches, allowing these clusters to collapse into sinking plumes. The resulting collective flow not only transports populations to depth without the energetic cost of self-propulsion but also enhances resource encounter through diffusivity-dependent molecular trapping. Via bioconvection, a concentrated patch of cells sinks under gravity, and those downwelling population migrate to depths without spending energy via active propulsion. Upwelling cells get aided by the flow fields they collectively self-generate, thus reducing energetic costs associated with vertical migration by orders of magnitude (See supplementary information, Fig. S11). Thus, age-dependent phenotypic changes influence the onset of bioconvection, promoting enhanced nutrient transport and mixing, thereby ensuring nutrient availability when needed the most.

Using microparticles as passive tracers (Fig. 2a, S4), we extend this understanding to ecological settings^57^, demonstrating that non-motile pico-plankton can exploit bioconvection to access nutrient-rich microenvironments that would otherwise remain inaccessible. Similarly, larger particles comparable to size of individual cell, representing dead or stressed cells of the same species, were passively entrained within bioconvective flows. We demonstrate the ecological relevance of self-generated bioconvective flows, by assessing the cell physiology under bloom-like settings. Ocean surface warming and stratification due to climate change promote algal bloom formation and can rapidly deplete nutrients in surface waters. Dense microbial accumulation formed during algal blooms can result in self-shading effect^6^, thereby reducing light availability across depths and posing a challenge for other photosynthetic species to thrive. Nutrient availability and acquisition get suppressed for blooming species, on top surface layers, owing to the density stratification. While stratification creates density barriers which suppress vertical nutrient exchange^47,48^, bioconvection may act as a collective mechanism that redistributes the local molecular landscape, reshaping the physical and chemical niche experienced algae under such constraints.

Under nutrient starvation, cells accumulate intracellular lipid droplets, which alter cell biomechanics and reduced migration; upon nutrient re-supply, these lipids disintegrate and phenotypic traits are recovered^56^. Bioconvection can significantly enhance the transport across density barriers, promoting rapid physiological recovery of the cells by improving nutrient access, consistent with the observed breakdown of intracellular lipids and the rapid restoration of photosynthetic efficiency^56^. Additionally, we observe that the mean swimming speed of the population recovered from 27 µm/s to 36 µm/s within 6 h, consistent with the faster nutrient replenishment mediated by bioconvective transport. This suggests that self-generated bioconvective flows not only drive active transport but also feedback on the cellular physiology, aiding motility and photo-physiological recuperation in ecologically meaningful ways. In natural settings, wind-driven shear can further enhance vertical exchange of dissolved molecules across stratified layers, potentially assisting the nucleation and onset of bioconvection plumes. More broadly, bioconvective dynamics emerging from competing, e.g., due to light and oxygen gradients in natural settings^33^, can govern multiscale active transport, mitigating resource limitations and enhancing microbial resilience across both marine and freshwater ecosystems.

