## Supplementary Materials for "Multiscale active transport driven by gravitactic bioconvection promotes resilience in algal blooms"

### This file includes:

Materials and methods

Tables T1-T2

Figures and captions S1 to S11

Supplementary Movies 1 to 11

References

1. Supplementary Discussion

i) Materials and methods
ii) Statistical tests
iii) Energy expenditure by motile cells in nutrient-limited environments: comparative estimation of the energetic costs for actively swimming gravitactic cells, versus collective cell migration due to self-generated bioconvection

2. Supplementary Tables

i) Supplementary Table 1: Glossary of symbols used in this study
ii) Supplementary Table 2: Age-dependent phenotypic variations in the cell cultures

3. Supplementary Figures

i) Fig. S1: Dynamical cell concentration and geometry of bioconvection
ii) Fig. S2: Orientational stability of *Heterosigma akashiwo HA3374*
iii) Fig. S3: Age-dependent variation of swimming stability within cell populations
iv) Fig. S4: Enhanced micro-cargo transport by bioconvective flows
v) Fig. S5: The distribution of cell traits determines the emergent range of hydrodynamic transport
vi) Fig. S6: Emergent offset of flow fields and finite-time Lyapunov exponents (*FTLE*s) in bioconvective flows
vii) Fig. S7: Molecular transport due to bioconvection
viii) Fig. S8: Bioconvective transport in a stratified environment accelerates nutrient supplementation and physiological recovery of cells
ix) Fig. S9: Bioconvective mixing facilitates nutrient transport across density interfaces, suppressing lipogenesis
x) Fig. S10: *NPQ* distribution for cell samples at different times, under nutrient supplementation
xi) Fig. S11: Schematic illustration of the geometric path length traversed by gyrotactic cells hitchhiking bioconvective flows

4. Supplementary Movies

i) Supplementary Movie 1: Onset of bioconvective plumes from a homogeneous suspension of motile algae
ii) Supplementary Movie 2: Cellular phenotypic traits underpin the onset of bioconvective patterns
iii) Supplementary Movie 3: Measuring orientational stability of motile phytoplankton
iv) Supplementary Movie 4: Cellular turnover time within bioconvecting plumes
v) Supplementary Movie 5: Quantifying cell-driven hydrodynamic fields through deep learning-based image segmentation

vi) Supplementary Movie 6: Simultaneous tracking of active and hydrodynamic flow fields in bioconvecting system
vii) Supplementary Movie 7: Single cell migrating pattern in self-generated flows
viii) Supplementary Movie 8: Molecular trapping and transport in bioconvective flows
ix) Supplementary Movie 9: Trapping transport contour via finite time Lyapunov exponent (*FTLE*) analysis
x) Supplementary Movie 10: Data-based simulation of molecular transport in bioconvective flows (case 1)
xi) Supplementary Movie 11: Data-based simulation of molecular transport in bioconvective flows (case 2)

5. References

### Materials and methods

#### Cell culture

The raphidophyte *Heterosigma akashiwo* (CCMP3374) was cultivated in sterile 50 ml glass tubes, exposed to a 14:10 h light-dark cycle to simulate natural diel rhythms within an incubator. The cells were cultured in f/2-Si medium (without silica) at a salinity of 36 PSU and a constant temperature of 22°C. During the illuminated phase, a light intensity of 1.35 mW was provided to ensure uniform exposure. Cell cultures were propagated by extracting 2 ml of suspension from 1 mm on the surface of the mother culture and transferring it into 25 ml of fresh medium every three weeks. Culturing of cells were carried out under sterile conditions within a laminar flow hood. The incubator was equipped with a white light source (λ = 535 nm, 1.35 mW) to replicate daylight conditions. All experiments were conducted between 12:00 and 16:00 h, a period during which a significant fraction of the cells exhibited negative gravitactic migration and bioconvection (Fig. S1. e), i.e., actively swimming upward against gravity. Within a millifluidic chamber (10 mm × 3.3 mm × 2 mm), we observed cell accumulation over growth phases of culture and found that bioconvection was absent before 240 h of post-inoculation. Bioconvection emerges in cell cultures spanning 360 h to 720 h after inoculation: an optimal time period for plume onset and stability. All experiments were carried out during this timeframe, and analysed for bioconvection with respect to the changing cell phenotypes.

#### Quantifying growth curve

To quantify the growth curve, fresh cultures were prepared, and measurements were carried out every 24 h. A volume of 5 µl was extracted from a uniformly homogenized culture tube and introduced into a small millifluidic chamber. After 144 h of post-inoculation, we dilute the sample volume with the culture supernatant of the same age, by half of the initial concentration, to ease image processing and post analysis in cell identification. Images were captured using stereo microscope (Nikon ® SMZ1270) for 10 s at a rate of 45 frames per second (fps). The identification of individual cells within these sequences of frames was extracted by image processing tool - OpenCV module in Python, employing size-based thresholding techniques. The counts obtained from this process were then averaged across all frames captured, with subsequent conversion into appropriate units across two biological replicates, with each having two technical replicates (four replicates in total). The experimental data follows a logistic fit (Fig. 1a) of the functional form:

$$f(x) = \frac{L}{(1 + e^{-k\,(x - x_0)})} - De^{-m(x - x_1)}$$

Where *L*: carrying capacity, *k*: growth rate, *x0*: midpoint of growth, *D*: depth of death phase, *m*: rate of decay, and *x1*: onset of death phase. The cell doubling time was obtained (Fig. 1a top inset) from the semi log linear fit at the exponential phase,

$$T_D = \frac{t \ln(2)}{\ln(n_f/n_i)}$$

where *t* is the time interval between initial and final measurement, $n_i$: initial cell concentration, $n_f$: final cell concentration.

**Cell motility estimation from single cell tracking**

We measure the swimming speed of cells within a custom-built millifluidic chamber made up of polymethyl methacrylate, measuring 10 mm (long) x 3.3 mm (high) x 2 mm (deep), attached to a *xyz* - translational cage. Before each experiment, we gently mixed the culture to ensure even distribution, then carefully take cell samples from the culture tube and injected them into the chamber. The chamber was placed on a stage that could move in three directions using micrometer screws. To observe the motility, we first let the millifluidic chamber, now filled with cells, reach stationary distribution, typically within 5 minutes. Then, an automated motor quickly flipped the chamber (Fig. S2 a) upside down by 180° in 3 s, thus changing cell orientation instantaneously with respect to the gravity vector. Cells undergo gravitactic reorientation (Fig. S2 b) and then migrate straight opposite to the direction of gravity. Their vertical movement was recorded for 40 s at 16 fps using a high-resolution camera and zoom lens (Navitar 6.5× Ultra Zoom, model 1-60191 AD, 12 mm FF) - coupled with a digital camera (Imaging Source DFK33UX265). A red LED (630 nm) was used for uniform and consistent lighting. To analyze movement, we used Python-based image processing (OpenCV2) to track individual cells (Fig. S2 C). The TrackPy module linked positions across frames, allowing us to measure the *x* (horizontal) and *y* (vertical) components of swimming speed (Fig. S5 c) in 4 replicates (2 biological with each having 2 technical replicates).

**Experimental setup and analysis of cell's reorientation time**

To measure the reorientational stability, cells were filled in the rectangular milifluidic chamber ($10\ \mathrm{mm}\ \times 3.3\ \mathrm{mm}\ \times 2\ \mathrm{mm}$) and mounted to a custom-made holding cage in a vertical plane. The cage was attached to a stepper motor, enabling the entire assembly to rotate as needed (Fig. S2 a). The movements of the motor were controlled using a driver connected to an Arduino board and automated by programming to rotate the motor for a specified duration. A camera with a variable zoom lens system was assembled on the other end to capture cells rotational movement for a given flip. Since most of the measurements were done during or after the exponential phase cells are diluted with culture supernatant to ensure better single cell tracking. We conducted two biological replicates with each of the replicates having three technical replicas (total 6 replicates), for each of the time point of observation. After assembling the millifluidic chamber filled with cells in a vertical plane, we wait for 10 minutes, allowing cells to accumulate at the top of the chamber. Once the cells are distributed across the top surface, we applied a 180° flip was given to the chamber. After the cells swim to around the middle of the chamber, the chamber was flipped again by 180°, allowing us to capture cell's reorientation tracks in the middle of the chamber (Supplementary Movie 3). Videos were acquired at a rate of 16 fps, and analysed using TrackPY python library. Trajectories, less than 45 continuous frames, were filtered out and the remaining tracks were interpolated (Fig. S2 c). To quantify cell stability, the rotation rate (ω) of cells was obtained as a function of their orientation relative to vertical direction for consecutive frames. For the given value of the angular position, the rotation rate was averaged (Fig. S2 d). The resulting distribution of ω (θ) was fitted with a sinusoidal function $A\sin(x+k)$, where *k* is the phase shift, related to cells rotational direction (Fig. S2 d, S3 a). The reorientation time scale was obtained from the amplitude of the best fit of sinusoidal function as $\tau_r = 1/2A$ [Suppl. Ref. 1]. To evaluate the distribution of reorientation across the population, the rotation rate and angular position of individual cells were tracked. The difference between the maximum and minimum values in the ω ~ θ distribution was identified as the stability parameter. The relative frequency distribution of this stability parameter was plotted (Fig. S3 c) for cells at different growth stages. To quantify the parameters (traits) most sensitive to age-dependent diversification, scatter distributions of motility and reorientation time were analyzed (Fig. S3 b). Correlation analysis revealed that reorientation time exhibited greater sensitivity to diversification as compared to motility, indicating increased heterogeneity in the population with respect to age (Fig. S3 b).

### Quantification of bioconvective flow by Particle Tracking Velocimetry (PTV)

#### i) The mixture of micro-particle tracers and cell culture

Polystyrene microparticles (PS), measuring 2 μm in diameter (SD = 0.04 μm), were selected as passive tracers to study emerging hydrodynamic environment due to bioconvection, by employing particle tracking velocimetry (PTV)s. A 100 μl volume of a PS solution, constituting a 10% w/v aqueous suspension, was sampled and subsequently diluted with 5 ml of MilliQ water. To achieve homogeneity, the mixed suspension underwent vortexing for a duration of 1 minute. Following this, a 1 ml sample from uniformly mixed solution was centrifuged at 3000 rpm for 5 minutes. For the experimental study on the trait-dependent bioconvective transport, we extracted the culture supernatant of the age, to dilute microparticle tracers. For extracting culture supernatant, we used a 0.2 μm pore-sized syringe filter (Whatman Puradisc 25 syringe filter). In the next step, 1ml of the extracted supernatant was employed to mix with the PS particles. The suspension of microparticle was mixed with the sampled cell culture, with 1:50 ratio. The resulting mixture was then carefully introduced into the millifluidic chamber.

#### ii) Image segmentation

To investigate the emerging hydrodynamics due to bioconvection, we used particle tracking when tracers are incorporated into the bioconvective system. We capture long frame sequences, extending up to a duration of 15 s at 45 fps. Image processing was performed through the openCV2 module within the Python environment. Individual cell coordinates were identified by employing binarisation and thresholding techniques (Fig. S4 a-c). The contours of both cells and particles annotated in the video data (Supplementary Movie 5) were extracted. Segregation of cells and particles within each frame was achieved based on the size threshold, enclosed by the contours. For enhanced visualisation, cells and particles were distinctly annotated (Supplementary Movie 5,6). The locations of particles and cells in subsequent frames were mapped onto a blank frame. This mapping allowed for an view of the fate and trajectories of particles (or cells) solely. Via image processing, we segregate the raw data having both cells and tracers into two distinct dynamic fields - i) hydrodynamic flow field, consisting movement of tracers only and ii) Bioconvection-driven cell's active flow field consisting movement of only the cells (Supplementary Movie 6).

### Mean squared displacement (MSD) estimation

We characterise the nature of the transport by quantifying the mean squared displacements (MSDs) of the particles advected by bioconvective flows. As a first step, we develop a pipeline for particle tracking (Fig. S4 a-c) and then, extraction of the trajectories (Fig. S4 d), and finally, the calculation of the mean squared displacements (Fig. S4 e). The extraction of trajectories was performed by linking particle co-ordinates using Track-Py module. Post-linking, we filter out the tracks exhibiting inconsistency over a span of 100 frames. To quantify the dynamic behavior of tracers, we computed the mean squared displacement of every single particle (after filtering out shorter tracks) over a temporal span of at least 100 frames. The ensemble mean of the MSD over time, was computed to extract coefficients and to analyze transport behavior of micro-cargo (Fig. S4 e, g). As a control reference, we conducted an experiment capturing the motion of particle in a quiescent fluid environment. Subsequently the log-log representation of MSD plots (Fig. S4 g) was fitted with formulated equation log (MSD) = $w_0 + w_1 \log(t)$, where the exponent $w_1$ (slope) provides information about the ballisticity inherent in the active transport of particles.

### Particle image velocimetry (PIV) analysis

For quantifying self-generated flow fields, Particle Image Velocimetry (PIV) analysis was carried out on the acquired movies, focusing on bioconvections with a single stable plume (Fig. S5 a). The preprocessing and velocimetry analysis were conducted using PIV lab (MATLAB), to extract the flow field information (*x, y, u, v*) and the further post processing and plots were done via Python code. The fast Fourier transform cross-correlation algorithm with 2-pass interrogation (128px, 64px) was implemented for this analysis in PIV lab. After the PIV analysis, post-processing was performed to filter out anomalous vectors, employing standard deviation and local median filter threshold. To quantify the range of active transport, we performed PIV analysis only on the hydrodynamic flow field. First, we perform image segmentation to identify tracers in bioconvective flows, and then the co-ordinate information of the tracers over time was mapped to an empty background, to highlight the motion of passive tracers. We employ PIV analysis on the tracer's motion data, using which the strength of the flow field was assessed, and a threshold streamline boundary was obtained in post-processing using Python. The active transport range threshold was estimated from the ratio of MSD exponent for the control scenario and at the plume center. Far from the plume, the transport exhibited diffusive (like in the control case) behavior and the ratio field strength are less than 60% of the field strength at the center (Fig. S5 b). This threshold provided a quantitative measure of the spatial extent of bioconvection-driven transport.

### Calculation of turnover timescale

To quantify the turnover timescale of cells hitchhiking in a bioconvective flow, the PIV data was obtained across different growth stages, with three biological replicates each. We perform numerical simulations from the extracted flow fields, with a sample size of 10 cells getting advected by the bioconvective flow. Let $t_k$ and $t_{k+1}$ be the times of two successive crossings with the same orientation, the turnover time is given by:

$$T_{to} = t_{k+1} - t_k$$

Passive particle trajectories were computed in a Lagrangian framework by integrating the advection equation given by,

$$\frac{d\mathbf{x}}{dt} = \mathbf{u}(\mathbf{x}, t),$$

$$\mathbf{X} = (\boldsymbol{x}, \boldsymbol{y}), \mathbf{U} = (\boldsymbol{u}, \boldsymbol{v})$$

Via code, we track the sign of $S(t) = x(t) - x_{in}$ ($x_{in}$ is the initial location of a trajectory) and register events where *S(t)* changes sign; the first pair of same-direction crossings separated by more than a small threshold defines the turnover time $T_{to}$. A trajectory is considered "closed" if such a pair is observed before the maximum integration time. The time required for these cells to complete a fully closed loop and return to their initial positions within a given tolerance was computed (Fig. S1 c, Supplementary Movie 4).

**Lyapunov exponent of the flow fields**

PIV-Lab was used to capture the velocity field of the stable bioconvective plume. The exported velocity field data was used for further analysis in Python. We used bilinear interpolation method that provides a continuous representation of the velocity filed over the specified domain of interest.

The finite-time Lyapunov exponent (*FTLE*) represents the finite-time average of the maximum expansion rate experienced by a pair of particles as they are transported within a fluid flow. We employed the existing method to compute *FTLE* and Lagrangian coherent structures (*LCS)* for the emergent flow filed in our system [2-4]. Given a velocity field *U(x,t)* defined over the spatial co-ordinates *x*, and time interval [*0, t*] the flow map $\varphi_t(x)$ is governed by the equation

$$\frac{d}{dt}\varphi_t(x) = \mathrm{U}(\varphi_t(x), \mathrm{t})$$

The equation describes the mapping of fluid particles over time due to the flow (Fig. S6 b).

In the next step the computation of the Jacobian matrix involves obtaining the spatial gradient of the flow map.

$$D\varphi_t(0,T) = \begin{bmatrix} \frac{\Delta x(T)}{\Delta x(0)} & \frac{\Delta x(T)}{\Delta y(0)} \\ \frac{\Delta y(T)}{\Delta x(0)} & \frac{\Delta y(T)}{\Delta y(0)} \end{bmatrix}$$

$$\sigma(x) = \frac{1}{T} \log \sqrt{\lambda \max{(\mathit{\Delta})}}$$

where $\mathit{\Delta} = [D\varphi_t(0,T)]^* \, [D\varphi_t(0,T)]$

" * " denotes the transpose of a matrix. Here, *σ(x)* is the *FTLE* at a given spatial point *x* is computed as the maximum eigenvalue over the specified time interval. Eigenvalues of the strain tensor are computed to quantify the stretching and compressing behavior of material elements. The ridge in the finite-time Lyapunov exponent (*FTLE*) field, identified as 'Lagrangian coherent structures (*LCS*)', (Fig. S6 c, Supplementary Movie 9) acts as a boundary with the highest local rate of separating nearby fluid trajectories. [4].

**Stratified environment in-a-chip**

To mimic stratification in aquatic ecosystems, a fluid column was prepared with density differences due to salinity. In natural settings, deeper layers are more saline and denser than the upper layers, creating a density barrier that hinders nutrient upwelling by diffusion. The objective was to investigate the feeding mechanisms of phytoplankton considering their collective scale emergent transport, when their population is trapped in the upper layer, specifically over shorter timescales compared to diel vertical migration (DVM). Typically, the culture medium has a salinity of 36 PSU, and at later saturation phases, it becomes nutrient-depleted. During the experimental phase, cells were in a nutrient-starved state. Two experimental conditions were established. A highly saline medium (42 PSU) containing essential nutrients (nitrate, phosphate, and other minerals) was prepared and mixed with a food dye to visualize molecular transport (Fig. S8 a). In the first setup (Experiment), 15 ml of 42 PSU medium was placed at the bottom, and 10 ml of highly concentrated starved cell culture (36 PSU) was gently introduced on top via a syringe pump. The high cell concentration generated bioconvection but barely penetrated the density barrier. In the second setup (Control), a similar stratification was created, but a diluted cell culture was placed on top to prevent convective plumes, allowing nutrient mixing to occur solely through diffusion. Over time, we compared the mixing across depths, between two cases via assessing the dissipation of chromatic dye (Fig. S8 b). samples were collected from the upper layer (Fig. S8 c), and lipolysis and photo physiology were analyzed (Fig. S8 d,e). The comparison between these setups demonstrated that in bioconvecting samples, active flows facilitated faster nutrient transport from deeper layers, whereas in the diffusive setup, nutrient supplementation and physiological recovery occurred at a slower rate. The physiological recovery were tracked via analyzing cell photo-physiology and lipolysis.

**Measuring photophysiology by pulse-amplitude modulated chlorophyll fluorometry (PAM)**

We used pulse amplitude modulated chlorophyll fluorometry (PAM) to quantify photosynthetic efficiency of *H. akashiwo* cells for two experiments with faster and relatively slower nutrient replenishing conditions over regular time intervals. We used a multiple excitation wavelength chlorophyll fluorometer (Multi-Color-PAM; Heinz Walz GmbH, Effeltrich, Germany) to quantify the maximum photosynthetic quantum yield ($F_v/F_m$), the maximum electron transport rate ($ETR_{max}$) and non-photochemical quenching (*NPQ*) of cell culture which is a measure of photo physiological status [5]. 1.5 ml of cell suspension sampled from the top of culture tubes for experiment and control conditions and filled into a quartz – silica cuvette (Hellma absorption cuvettes; spectral range, 200 to 2500 nm; pathlength, 10 mm). The samples were dark adapted for 5-7 minutes before the initiation of measurements [6]. We used Multi-Color PAM 3 Win software saturation pulse (SP) and light curve method at specified time intervals to quantify $F_v/F_m$, and $ETR_{max}$ and for both the control (C) and experimental (E) case.

**Quantification of intracellular lipid area**

To quantify the size of lipid droplets inside the cell body, sample solutions from the culture tubes were stained with Nile Red (Thermo Fisher Scientific; excitation/emission, 552/636 nm). We dissolved 10 µl of Nile red + dimethyl sulfoxide (DMSO) solution to 200 microliters of culture supernatant and thoroughly mixed. The resultant solution is again mixed with 200 µl of cells and incubated in dark for 10 minutes. We used fluorescence microscopy (Olympus CKX53 inverted microscope, Imaging Source, DFK33UX265) to identify and characterize the accumulation of LDs in single cells (Fig. S9). Via image processing we extracted the cell contour area, and the area bounded by all lipid droplets inside the cells via thresholding. The sum of each of the lipid contour areas is then normalized with cell area to obtain the required parameter. (Fig. S8 d).

**Simulation of molecular transport due to bioconvection**

On obtaining the time-varying velocity field from the PIV data, a time-averaged velocity field is constructed, which served as the underlying velocity field for the advection of the tracer. The time-averaged velocity field is simply found using the form

$$\langle u_i \rangle = \frac{\sum_{j=1}^{N} u_i}{N},$$

where $u_i$ denotes a vector component of the velocity field and *N* denotes the number of PIV time frames while the time averaged operator is represented by $\langle\ \rangle$. With this background velocity field, a time-dependent convection-diffusion equation is solved having the form

$$\frac{\partial c}{\partial t} + \langle \boldsymbol{u} \rangle \cdot \nabla c = D \nabla^2 c$$

where *c* is the molecular concentration and *D* is the diffusivity (in $m^2/s$). Here $\langle \boldsymbol{u} \rangle = \langle u_i \rangle$ represents the time-averaged velocity field vector. Different initial conditions and magnitudes of diffusivity influence the evolution rate and fate of the molecular concentration field. To this effect, *D* is varied between $5 \times 10^{-8}$ and $5 \times 10^{-11}$ $m^2/s$. The varying location of molecular introduction (varying initial conditions) are discussed in the main text wherein we note that the concentration ($c$) is 1 at the location where it is introduced while the rest of the domain has $c$ = 0. The domain for the simulation is the same as obtained from the PIV results. The top boundary is assumed to be impermeable and thus a no-flux boundary condition $\boldsymbol{n} \cdot [-D\nabla c + \langle \boldsymbol{u} \rangle c] = 0$ is employed in the top wall. The lateral walls are assumed to have a Direchlet boundary condition $[c = 0]$ to mimic the far-stream situation where the concentration of the tracer is negligible. The bottom boundary may have either a no-flux condition (implying an impermeable wall) or a no-diffusive-flux condition $[\boldsymbol{n} \cdot D\nabla c]$ (implying that any evolution of the tracer concentration across that boundary is driven by advection). Here $\boldsymbol{n}$ denotes the outward normal from the boundary.

The mixing of the molecules is analyzed employing the scalar dissipation rate parameter ($\chi$), which is defined as

$$\chi = 2D(\nabla c)^2$$

Averaged scalar dissipation rate can be obtained as $\langle \chi \rangle = \int_{\partial \Omega} \chi dA \big/ A$. $\langle \chi \rangle$ delineates the mixing possibility of a system. A higher value of $\langle \chi \rangle$ implies large concentration gradients, thereby, shows that the system possesses a high mixing capability. In contrast, a low value of $\langle \chi \rangle$ implies good mixing has taken place. A rapid decrease of $\langle \chi \rangle$ with time implies the system is undergoing excellent mixing of the tracer, while an increase in $\langle \chi \rangle$ with time shows that the background velocity is effective de-mixing or concentrating the tracer sample. We note that the unit of concentration can be arbitrary in the time-dependent convection-diffusion equation, although usually the concentration unit is considered as $mol/m^3$. The above equation along with the appropriate boundary conditions is solved using the convection-diffusion model in the FEM package COMSOL Multiphysics. A free triangular mesh with maximum element size of $4.4 \times 10^{-5}$ is used for the study which results in $\sim 2 \times 10^4$ domain elements. The equation is solved till *t* = 1000 s using a fully coupled PARDISO solver.

### Statistical tests

A Welch's t-test was performed to compare the means of the two groups, accounting for unequal variances and sample sizes. The test statistic (t) was calculated as:

$$t = \frac{<X_1> - <X_2>}{\sqrt{\frac{S_1{}^2}{n_1} + \frac{S_2{}^2}{n_2}}}$$

where $< X_1 >$ and $< X_2 >$ represent the sample means of the two groups, $S_1$ and $S_2$ are the standard deviations, and $n_1$ and $n_2$ denote the sample sizes.

The degrees of freedom (*df*) were calculated using the Welch-Satterthwaite equation:

$$df = \frac{(\frac{S_1{}^2}{n_1} + \frac{S_2{}^2}{n_2})^2}{\frac{(\frac{S_1{}^2}{n_1})^2}{n_1 - 1} + \frac{(\frac{S_2{}^2}{n_2})^2}{n_2 - 1}}$$

A two-tailed test was conducted to determine whether there was a statistically significant difference between the means of the two groups. The corresponding p-value was obtained by computing the survival function of the t-distribution for the absolute value of the test statistic. Statistical significance was evaluated at a significance level (α) of 0.05 denoted by asterisks " * ". If the p-value was less than the significance threshold, the null hypothesis (H0) which states that there is no significant difference between the means was rejected in favor of the alternative hypothesis (HA), indicating a statistically significant difference between the two groups. The analysis was performed using Python's SciPy library.

### Energy expenditure by motile cells in nutrient-limited environments

At low Reynolds numbers, where viscous drag forces dominate, the energy required for phytoplankton to overcome drag is proportional to the viscosity of the surrounding medium and the velocity of swimming. The energy expenditure, $E_{cell}$, for a phytoplankton moving a given distance L is:

$$E_{cell} = F_d . L$$

where $F_d$ is the drag force acting on the cell is given by $F_d = 6\pi\eta V_{cell} r$ , for a spherical phytoplankton cell of radius $r$, swimming at velocity $V_{cell}$ in a fluid medium with dynamic viscosity $\eta$.

The distance $L$ can be traversed either by active propulsion alone or with assistance from bioconvective flow. In both cases, the energy spent is:

$$E_{cell} = F_d . V_{cell} . t$$

The key difference between these two scenarios lies in the timescale $t$, as $E_{cell} \, \alpha \, t$. In bioconvective flow, the turnover timescale is significantly faster, meaning that a cell spends less time actively propelling itself per cycle. As a result, the energy required for locomotion is reduced when assisted by bioconvection, enhancing overall efficiency in nutrient acquisition and transport.

*Case 1: Energy expenditure of a cell in actively propelling its body length through the geometrical path*

Phytoplankton rely on active propulsion to navigate through the water column, particularly in environments where fluid motion is limited. The energy expended by an individual swimming cell can be quantified using the expression

$$E_{cell} = F_d . L_{total}$$

$L_{total}$ is the total distance covered during its trajectory (schematic illustration – Fig. S11). The total path length comprises three main segments. The cell descends directly covering distance $L_{down}$ by collective sinking. Upon reaching a lower stratified or denser layer, the cell often undergoes a lateral shift $L_{near}$ before reorienting itself. After reorienting, the cell swims back towards the upper layers following $L_{arc}$. Thus, the total length of the trajectory can be expressed as $L_{total} = L_{down} + 2L_{near} + L_{arc}$. Using experimentally realizable approximations, the energy required for an individual cell to cover the total migration path under active propulsion can be rewritten as

$$E_{cell}(\text{swim}) = 6\pi\eta V_{cell} r\ (4+\pi)\ R$$

By substituting these values into the equation, we estimate the energy required for a single migration cycle through active propulsion without the assistance of ambient flow.

Assuming $R = 1$ mm, $V_{cell}$= 20 μm/s and $r = 10$ μm, the energetic cost of migration obtained to be $E_{cell}(\text{swim}) = 2.7 \times 10^{-14}$ J.

*Case 2: Energy Expenditure with Bioconvective Flow Feedback*

In a bioconvective environment, cells experience different energy expenditures depending on their position within the flow field. Unlike Case 1, where phytoplankton rely solely on active propulsion to traverse the water column, bioconvection introduces flow feedback mechanisms that assist in vertical and lateral transport. This significantly reduces the energy required for active swimming, as cells interact dynamically with the convective currents. The energy expenditure in bioconvection can be divided into distinct phases, corresponding to different regions of the migratory path.

**a: Passive sedimentation at the plume center**

When a cell enters a bioconvective plume, it experiences enhanced downward motion due to the bulk fluid movement. If positioned at the center of the plume, the cell predominantly sediments passively under gravity rather than expending energy for active propulsion. The energy required for this phase is negligible:

$E_{cell}(\text{BC}_{\text{center}}) = 0$

Since sedimentation occurs due to the downward convective flow, no additional energy input from the cell is needed during this phase.

**b: Reorientation with feedback from self-generated flow**

Once a cell reaches the lower region of the bioconvective flow, it must reorient and ascend back toward the upper layers. However, instead of relying entirely on active propulsion, the cell benefits from flow feedback. The total migration distance during reorientation, denoted as $L_{near}$, consists of both actively swimming and being advected by the bioconvective flow field.

To quantify the energy expenditure in this phase, we define an effective swimming distance $L_{eff}$, which represents the portion of $L_{near}$ where the cell actively propels itself. The energy required for reorientation within a convective flow field is given by:

$$E_{cell}(\mathrm{BC}_{\mathrm{Re0}}) = F_d.\ L_{eff}$$

The effective distance for active propulsion is expressed as: $L_{eff} = V_{cell}\ .\ t_{eff}$

Where $t_{eff}$ is the timescale for which a cell would spend its energy in swimming the distance $L_{eff}$. Since during this timescale the cell also experiences advection by the bioconvective flow, the effective length can be rewritten as

$$L_{eff} = V_{cell} \cdot \frac{L_{near}}{(V_{cell} + V_{adv})}$$

For cases where the advective velocity is significantly higher than the swimming velocity ($V_{adv} \gg V_{cell}$) the expression simplifies to: $L_{eff} \sim V_{cell}\ .\ \frac{L_{near}}{V_{adv}}$

$$E_{cell}(\mathrm{BC}_{\mathrm{Re0}}) = 6\pi\eta {V_{cell}}^2\ r\ .\ \frac{L_{near}}{V_{adv}}$$

$$E_{cell}(\mathrm{BC}_{\mathrm{Re0}}) = 6\pi\eta {V_{cell}}^2\ r.\ \frac{L_{near}.\rho H}{\eta.Re}$$

Where $Re =$ Reynold's number, $H =$ characteristic length scale and $\rho =$ density of fluid.

Approximating $L_{near} = R,\ H = 3R$

$$E_{cell}(BC_{ReO}) = 18\pi\eta {V_{cell}}^2 r\ .\ \frac{\mathrm{R}^2.\rho}{\eta.Re}$$

By substituting these values into the equation, we estimate the energy required while reorientation with feedback of bioconvective flow. ($Re$ ~ 10 for bioconvective flow)

$E_{cell}(BC(Reo)) =\ 2.26 \times 10^{-17}$J

**c: Propulsion with feedback from self-generated flow**

In this phase of bioconvective transport, a phytoplankton cell utilizes both its active propulsion and the upward convective currents generated by the collective movement of the population. Cells reaching the bottom of a bioconvective plume experience upwelling, where the convective flow assists in lifting them back toward the upper layers of the water column. However, active propulsion is still required for fine-scale control over positioning and reorientation. The total energy spent by a cell during this phase is expressed as:

$$E_{cell}(BC_{upwell}) = F_d\ .\ V_{cell}\ .\ t_{eff}$$

Since the cell is transported by the flow over the total distance of $L_{arc} + L_{near}$ , the effective time for which active propulsion contributes is given by

$$t_{eff} = \frac{(L_{near} + L_{arc})}{V_{adv}}$$

$$E_{cell}(BC_{upwell}) = 6\pi\eta {V_{cell}}^2 r\ \frac{(L_{near}+L_{arc}).\rho H}{\eta.Re}$$

$$E_{cell}(BC_{upwell}) = 6\pi\eta V_{cell}{}^{2} r \, \frac{(R+\pi R).\rho 3R}{\eta.Re}$$

$E_{cell}(BC_{upwell}) = 9.35 \times 10^{-17}$J

Addition of all the sub-cases results in the net energy spent by a single cell in bioconvective flow to propel the same geometrical distance.

$E_{cell}(BC_{total}) = 11.61 \times 10^{-17}$J,

which is three orders of less magnitude in comparison to energy spent by a swimming cell via active propulsion.

### TABLE T1. Glossary of symbols used in the study

| Physical and statistical parameters | Units | Symbol |
|---|---|---|
| Length of plume | m | $L_p$ |
| Width of plume | m | $W_p$ |
| Vortex offset | m | - |
| Vortex height | m | $Y_v$ |
| Hydrodynamic plume width | m | $W_{hyd}$ |
| Turnover time | s | $T_{to}$ |
| Depth averaged velocity | m/s | $\langle \lvert U \rvert \rangle_y$ |
| Mean squared displacement | $m^2$ | MSD |
| gravity | $m/s^2$ | g |
| Length of enhanced hydrodynamic transport | m | $L_{eh}$ |
| finite time Lyapunov exponent | 1/s | $\sigma_{ftle}$ |
| Scalar dissipation rate | 1/s | $\langle x \rangle$ |
| Molecular diffusivity | $m^2/s$ | $D$ |
| Aspect ratio | 1 | $AR$ |
| Probability density function | 1 | PDF |
| Velocity correlation | 1 | $V_{corr}$ |
| **Cell parameters** | **Units** | **Symbol** |
| Swimming speed | m/s | $V_{cell}$ |
| Vertical speed | m/s | $V_y$ |
| Rotation rate | rad/s | ω |
| Angular position | degree | θ |
| Reorientation time | s | $\tau_{re}$ |
| Stability parameter | 1/s | $A$ |
| Cell area | $m^2$ | $A_{Cell}$ |
| Lipid area | $m^2$ | $A_{Lipid}$ |
| (Relative) Electron transport rate of PSII | 1 | $ETR$ |
| Maximum quantum efficiency of PSII | 1 | $F_v/F_m$ |
| Non-photo-chemical quenching | 1 | $NPQ$ |
| Photosynthetically active radiation | μmol/m² s | PAR |

### TABLE T2. Age-dependent variations in cell cultures

| Age (hours) | Cell concentration ($\times 10^5$/ml) | Vertical speed (μm/s) | Re-orientation time (s) | Lipid area / Cell area |
|---|---|---|---|---|
| 360 h | $1.75 \pm 0.20$ | $65.0 \pm 10.9$ | $4.86 \pm 0.72$ | $0.06 \pm 0.02$ |
| 504 h | $1.87 \pm 0.31$ | $56.2 \pm 8.5$ | $5.55 \pm 0.77$ | $0.09 \pm 0.03$ |
| 672 h | $1.15 \pm 0.33$ | $52.8 \pm 18.1$ | $7.39 \pm 0.80$ | $0.12 \pm 0.03$ |

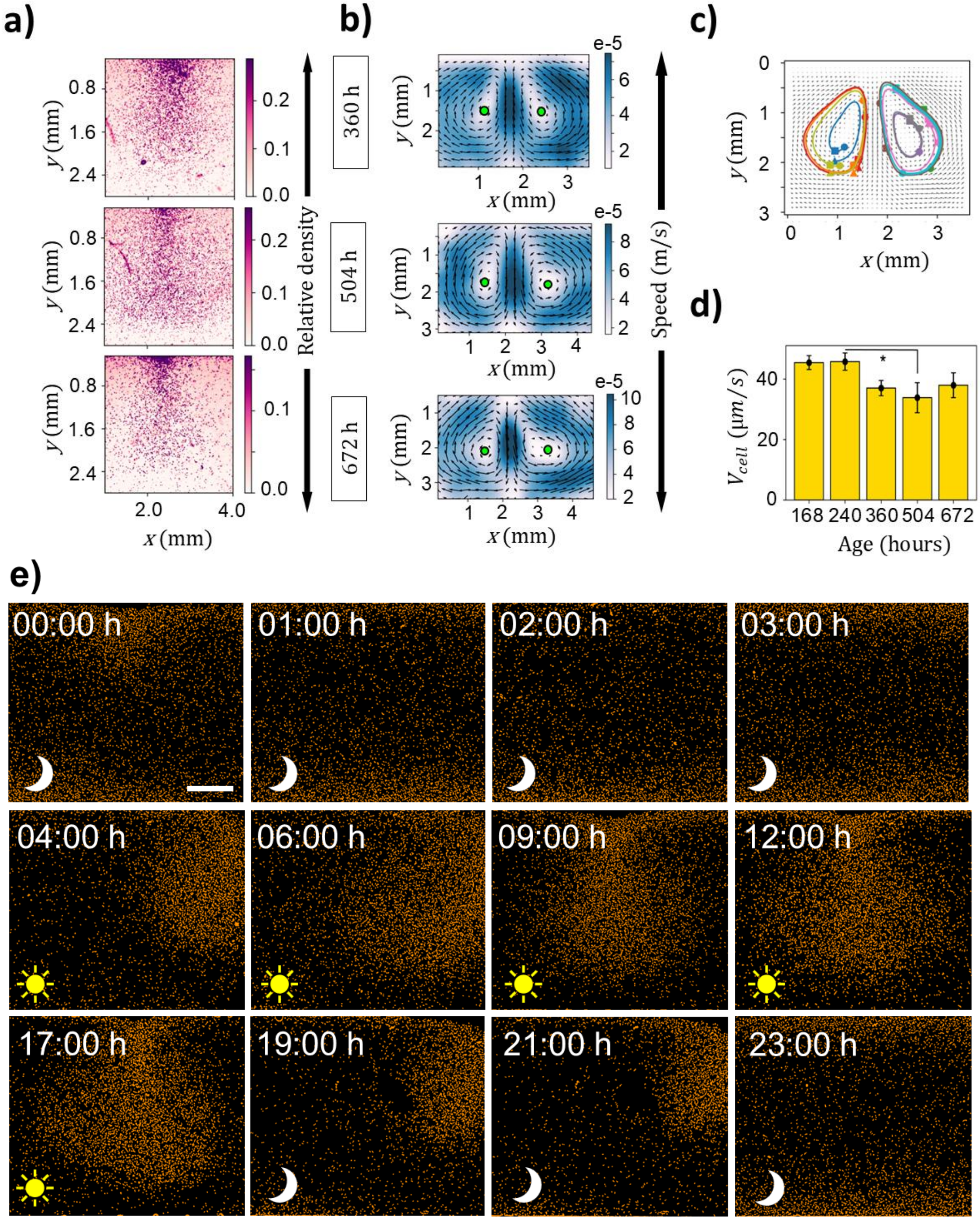
a)
b)
c)
d)
e)
Relative density
Speed (m/s)
360 h
504 h
672 h
e-5
$y$ (mm)
$x$ (mm)
$V_{cell}$ ($\mu m/s$)
Age (hours)
168 240 360 504 672
00:00 h
01:00 h
02:00 h
03:00 h
04:00 h
06:00 h
09:00 h
12:00 h
17:00 h
19:00 h
21:00 h
23:00 h

**Fig. S1: Dynamic cell concentration and geometry of bioconvection. a)** Local cell concentration (measured as mean relative density) in steady state plumes varies with age-dependent phenotypes (each row corresponds to a different cell age in hours). **b)** Velocity fields and vortex centers of bioconvective flows, showing age-dependent plume variation and downward shifts in vortex core position. **c)** Trajectories obtained from numerical computation to extract the time taken by cells complete one loop - turnover time from the PIV of cell's flow field (Supplementary Movie 4). **d)** cell swimming speed obtained from four biological replicates. Error bars represent mean $\pm$ 1.s.d. Asterisks indicate statistical difference (0.01< P <0.05) **e)** Bioconvection at different time points revealed relatively weaker flow dynamics during the night-time, due to bottom accumulation (scale bar – 0.5 mm). Cells were identified via image processing and annotated with color.

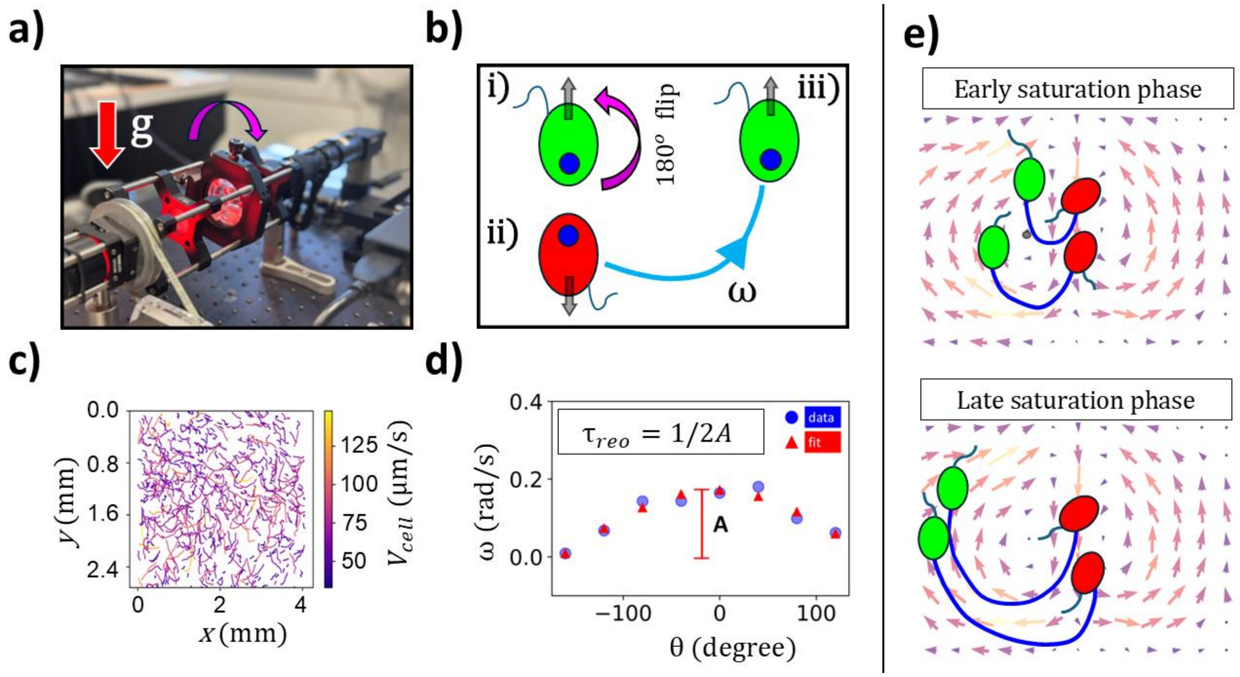


**Fig. S2: Orientational stability of *Heterosigma akashiwo HA3374.* a)** Experimental setup for imaging cell reorientation, featuring a programmable stepper motor connected to a flipping chamber to rapidly alter the cells' gravitactic orientation. **b)** (i) Negatively gravitactic species exhibit stable upward alignment. (ii) Upon 180° inversion of the chamber, cells lose alignment stability and iii) gradually reorient back to its stable position which is upward aligned. The angular speed of reorientation depends on the growth phase of the cells (Supplementary Movie 3). **c)** Trajectories of reorienting cells follow an arc-shaped path. **d)** Rotation rates extracted relative to angular positions along reorientation tracks, exhibit a sinusoidal pattern. Experimental data fitted with a sine curve, where the amplitude represents stability parameters. **e)** Schematics showing how collective reorientation shapes the geometry of steady state bioconvective flow. At late saturation stage, increased reorientation time leads to broader plume widths. As a measure of gravitactic sensing, reorientation time governs the ability of cells to escape the plume center early (or late) and maintain gyrotactic focusing along small (or large) closed-loop trajectories.

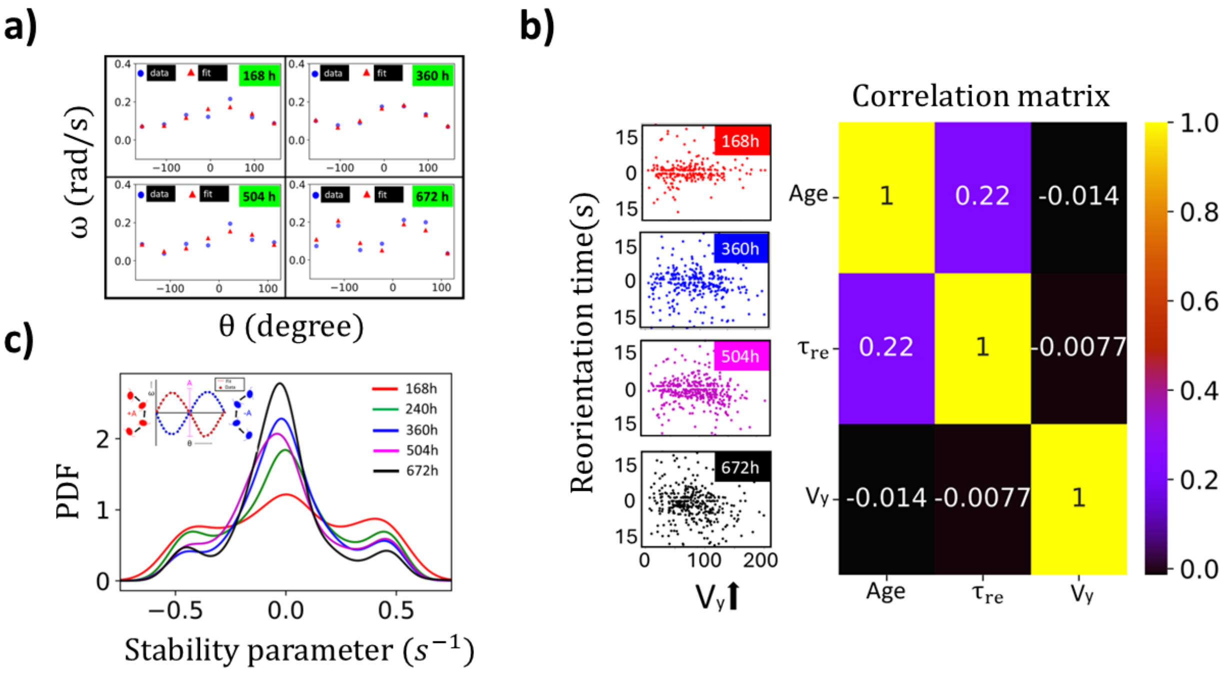


**Fig. S3: Age dependent variation in the stability parameter within population. a)** rotation rate obtained with respect to instantaneous angular position for cells of different growth stages. Scatter points in blue represent the experimental data with sine fit, represented as red scatter points. **b)** Reorientation time and upward velocity are uncorrelated, as shown by the random distribution of scatter points across growth phases. This suggests that reorientation is unbiased by vertical speed. Age-dependent reorientation adjusts more rapidly compared to age-dependent changes in speed. **c)** Probability distribution function of stability parameters within the population. The inset shows the clockwise and counterclockwise rotation patterns used by cells to stabilize their positions.

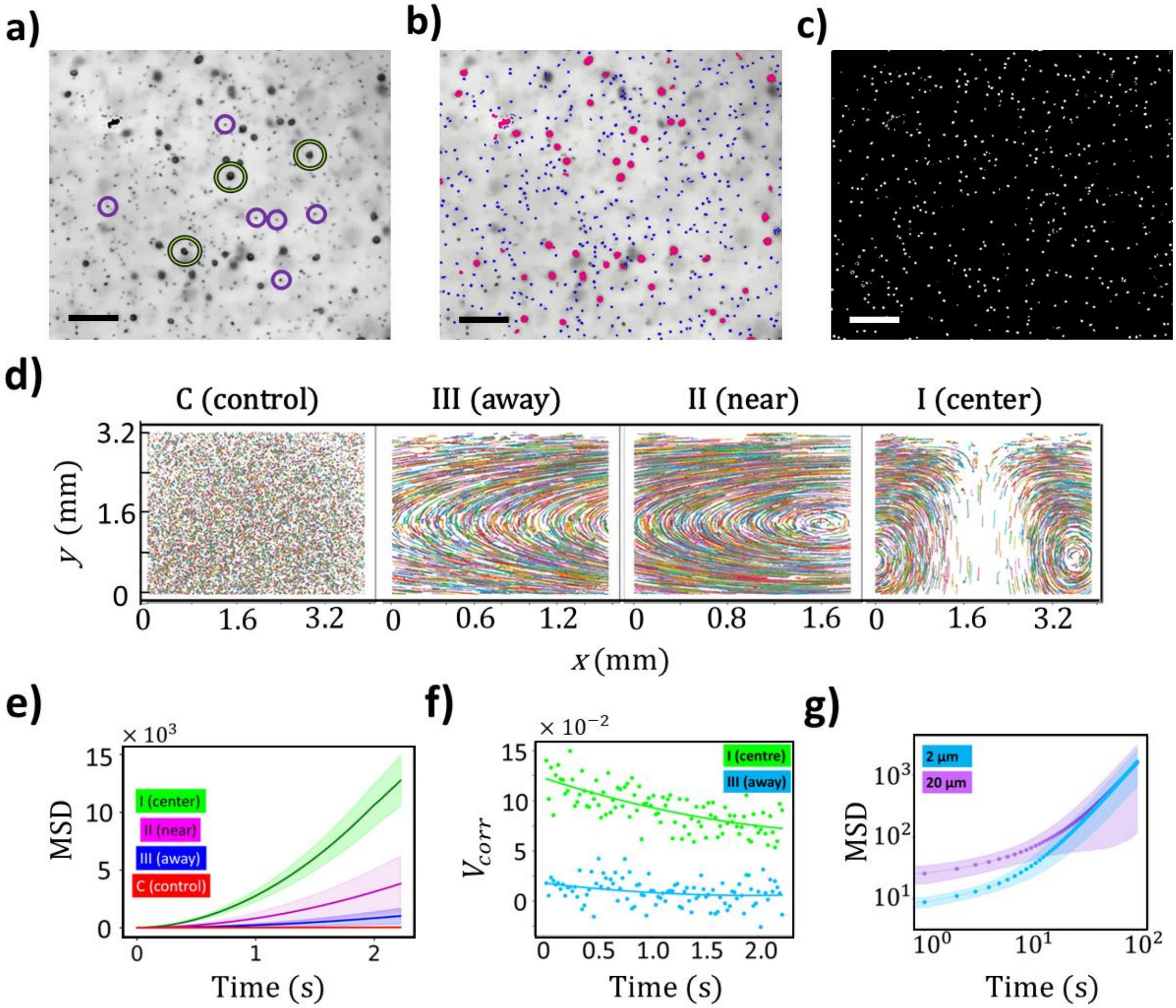


**Fig. S4: Enhanced micro-cargo transport by bioconvective flows. a, b, c)** Image analysis pipeline to perform particle tracking velocimetry. Image a show the captured frames with passive tracers incorporated in bioconvective flow (scale bar – 0.15 mm). Cells are comparatively bigger in size. **b)** By employing size-based filtering, image segmentation was done identifying cells and particles in the flow field (Supplementary Movie 5). **c)** The particle co-ordinates are mapped to a blank space to obtain the flow field of passive traces only (scale bar – 0.15 mm). **d)** Trajectories of neutral microparticle tracers advected by bioconvective flow at different regions: (I) plume center, (II) near the plume vortex—representing the collective upwelling of cells, and (III) away from the plume. The control condition represents passive particles floating in still fluid. **e)** Mean squared displacement (MSD) of particles in bioconvective flow across regions I, II, and III, compared to passive diffusion in still fluid. **f)** Velocity correlation plot indicating that passive tracer particles at the plume center exhibit highly directionally correlated motion. Particles away from the plume center show diffusive transport, hence the correlation between particles are comparatively lower. **g)** MSD analysis of two different particle sizes in bioconvective flow, showing that larger particles experience reduced ballistic transport, but still effectively captured in vortex rolling advection.

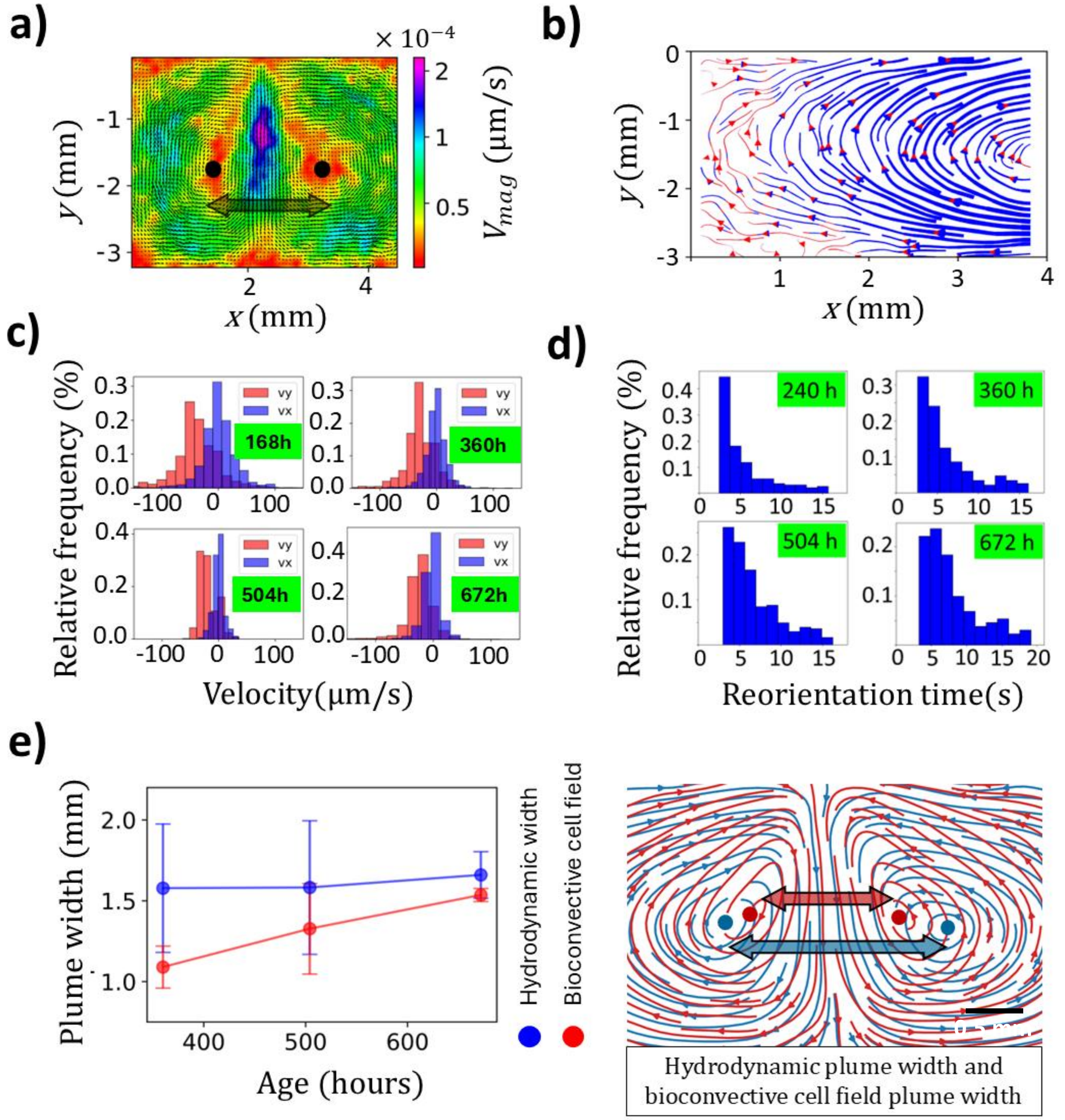


**Fig. S5: The distribution of cell traits determines emergent range of hydrodynamic transport.** **a)** Velocity field of bioconvective flow obtained through particle image velocimetry (PIV) analysis. **b)** Range of active transport within a bioconvective plume, shown by blue tracks indicating a threshold based on the strength of the actively driven flow field. Red tracks denote transport streamlines with a field strength reaching 60% of the central flow field strength. **c)** Distribution of the *x*- and *y*-components of the velocity field for cells across different growth phases. **d)** Relative distribution of cell reorientation times at different growth phases. **e)** Plume width of hydrodynamic and cell-movement fields in bioconvective flow (Supplementary Movie 6). For each flow field, plume width was calculated as the distance between two vortex centers, averaged over three replicates (scale bar – 0.5 mm).

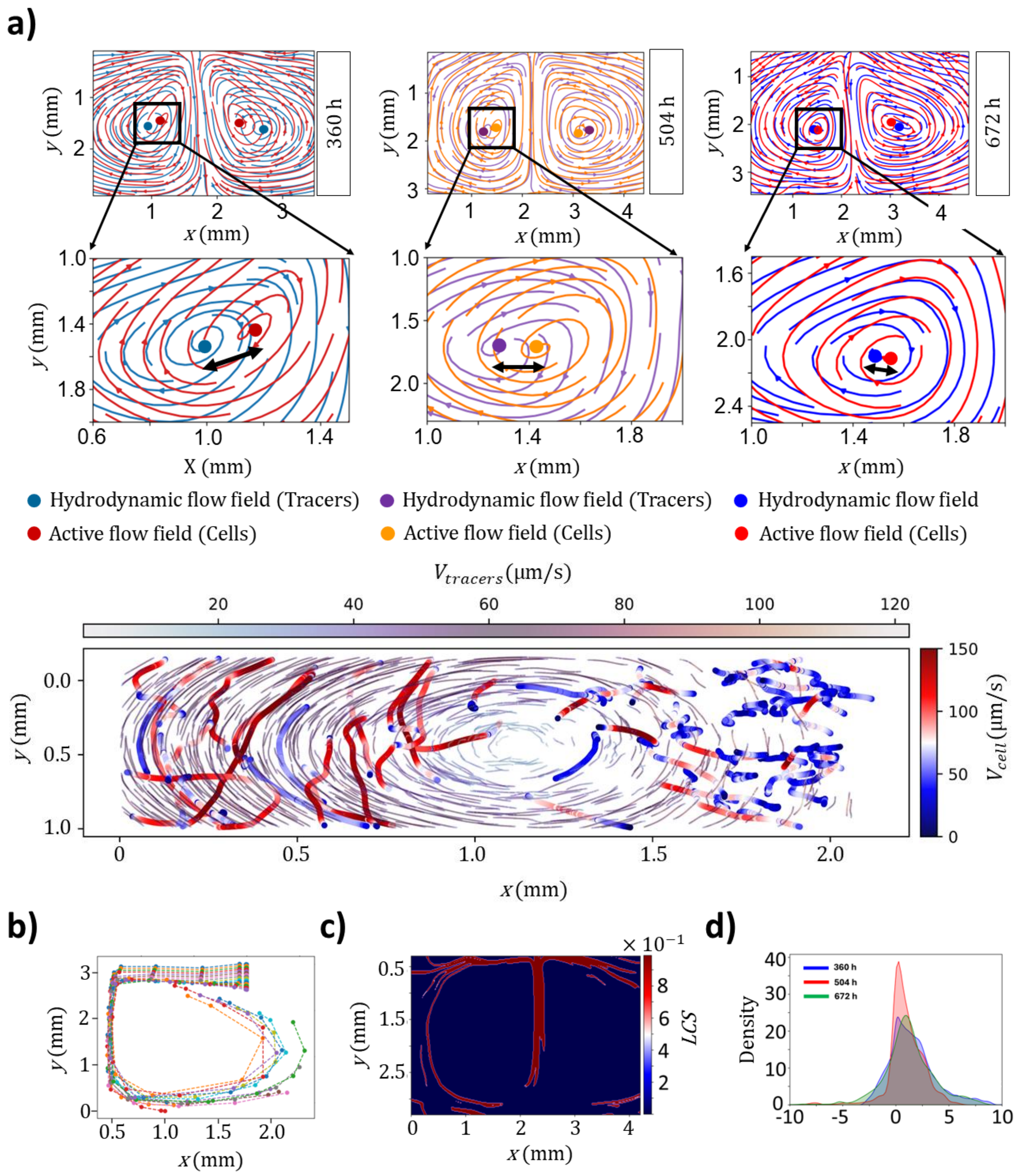


**Fig. S6: Emergent offset of flow fields and finite-time Lyapunov exponents (*FTLE*s) in bioconvective flows. a)** Streamlines of hydrodynamic fields and cell-movement fields in bioconvective flows across different generational timescale. The zoomed-in view shows that with aging, the vortex centers of the two fields coincide, indicating reduced cellular activity. Tracking of the cells and particles close to the vortex reveals mechanistic insight on these differences in flow fields (Supplementary Movie 7). Locations where the downwelling tracer speed opposes the upwelling gravitactic phytoplankton lead to short, confined cell tracks (around $x$ = 2mm). In contrast, where tracer and cell migration directions align, the tracks become long and persistent (around $x$ = 0). **b)** Flow map illustrating the advection of tracers initiated within the flow field. **c)** Lagrangian coherent structures (*LCS*) extracted from the flow field using the Jacobian transformation tensor (Supplementary Movie 9). **d)** *FTLE* distribution for an ensemble of particles over the simulation timescale, showing convective plume dynamics across different cell ages.

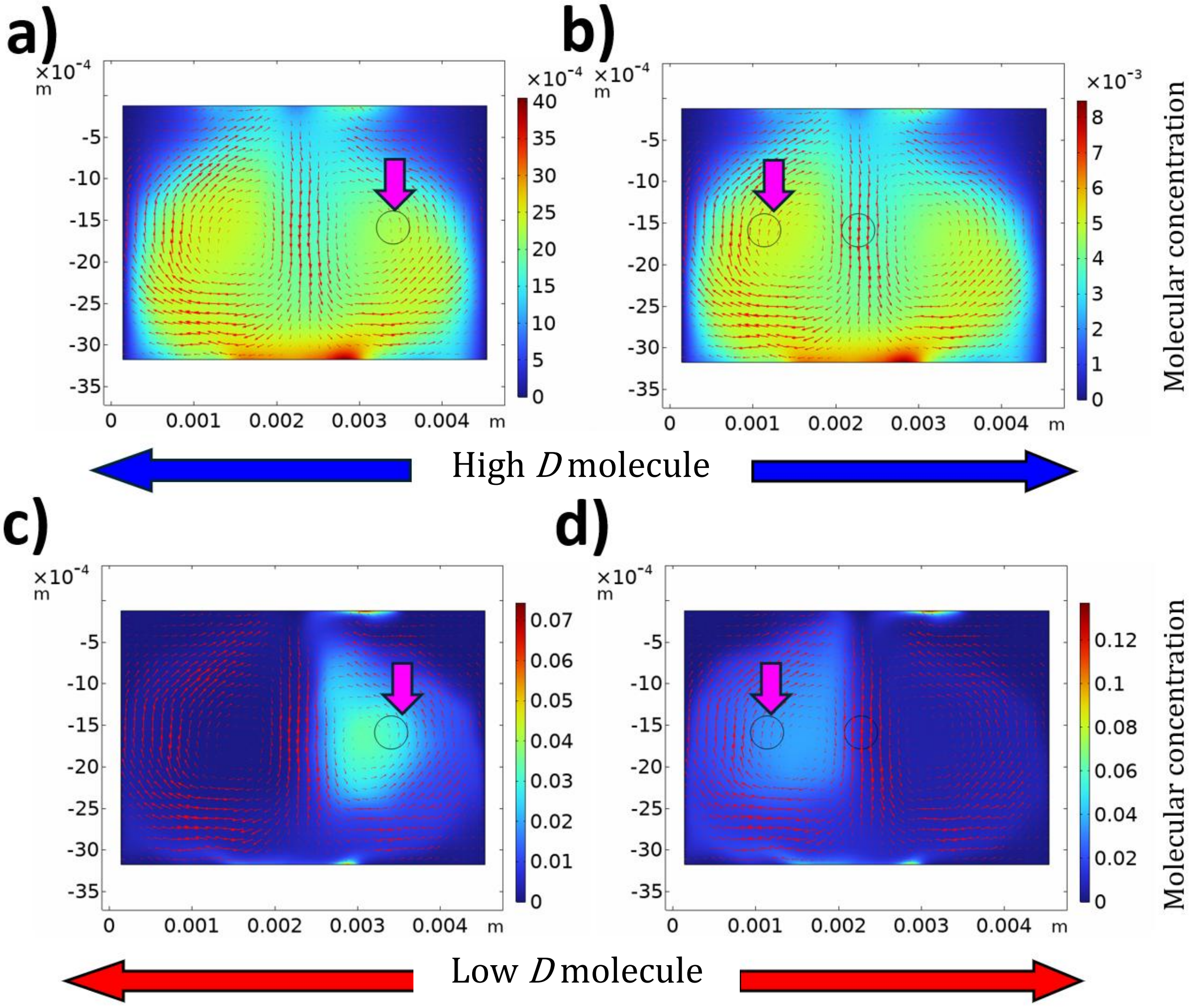


**Fig. S7: Molecular transport due to bioconvection. a,b)** Trapping of high-*D* molecules in the bioconvective flowe. Initial patches of molecules are released at the left and right vortex centers bioconvective flows **c,d)** Low-*D* molecules were not trapped, bur released into the environments, when initial patch of molecules are placed at the same vortex core.

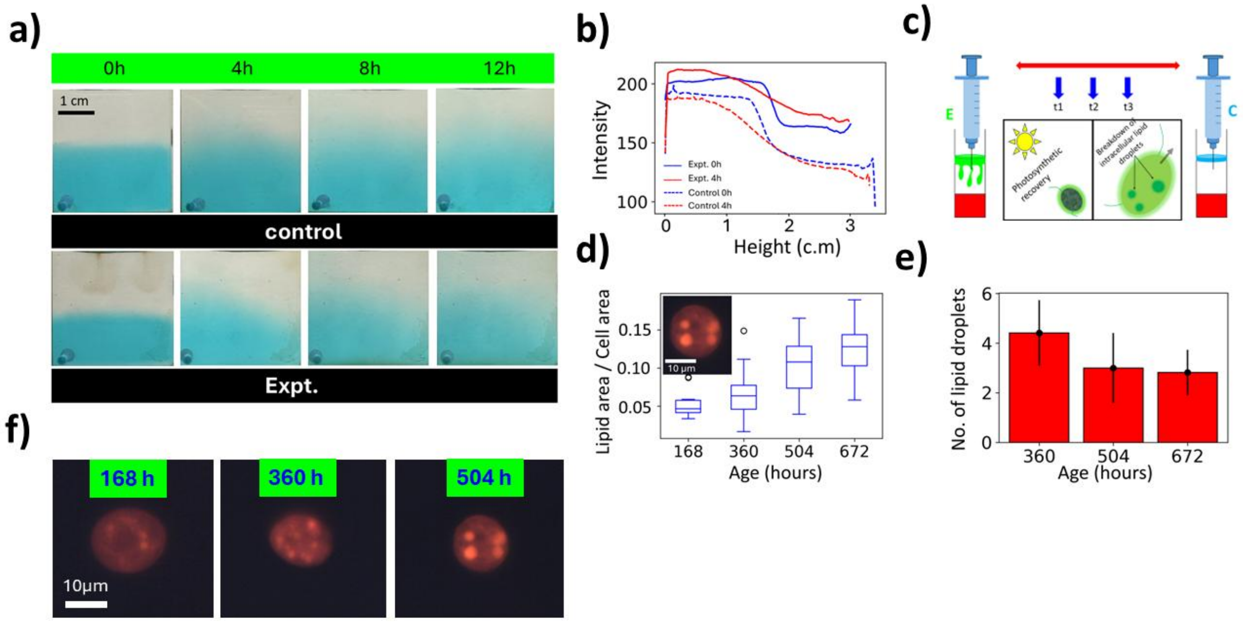


**Fig. S8: Bioconvective transport in a stratified environment accelerates nutrient supplementation and physiological recovery of cells**. **a)** Experimental setup showing a stratified system with a highly saline, nutrient-rich lower layer (colored with food dye) and a nutrient-absent upper layer. In the control, diffusion is the sole mechanism driving stratification breakdown and nutrient homogenization. In experimental conditions, bioconvective flow in the upper layer enhances stratification disruption. **b)** Mixing rate comparison over 4 h, where intensity stabilization indicates the homogenization of the colored dye throughout the column height. **c)** Schematic representation of collected samples from both experimental conditions, analyzed to assess physiological responses to nutrient supplementation. **d, f)** Lipid accumulation in cells as environmental nutrients become depleted. **e)** Over time, lipid droplets coalesce and merge into larger structures.

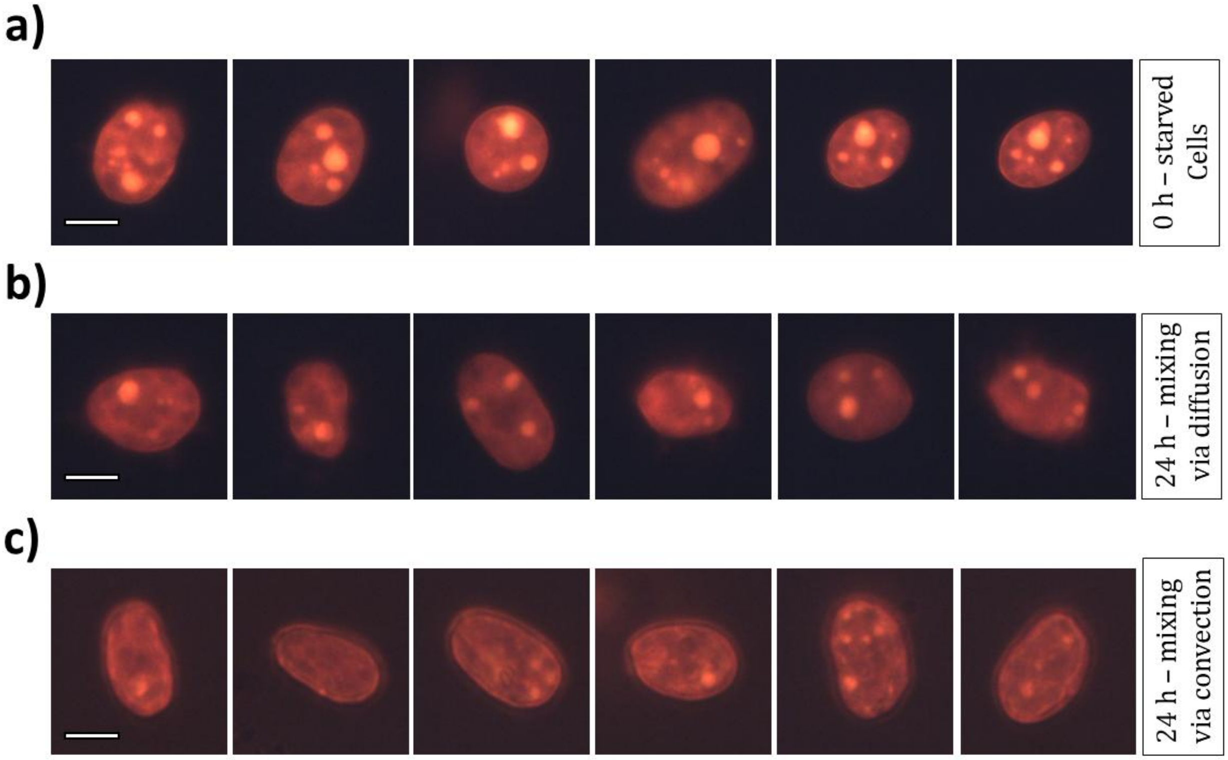


**Fig. S9: Bioconvective mixing facilitates nutrient transport across density interfaces counteracting lipogenesis. a)** Large and distinct lipid droplets in cells at 504 h under nutrient starvation pictured via staining with Nile red (scale bar – 7 µm). **b)** Control cells sampled after 24 hours (fig S8.c), from the stratified setup, where the nutrient supplementation is via diffusion (scale bar – 7 µm). **c)** Almost no lipid droplet formation for the cells sampled from experimental setup where the mixing process across the density barrier took place via bioconvection (scale bar – 7 µm).

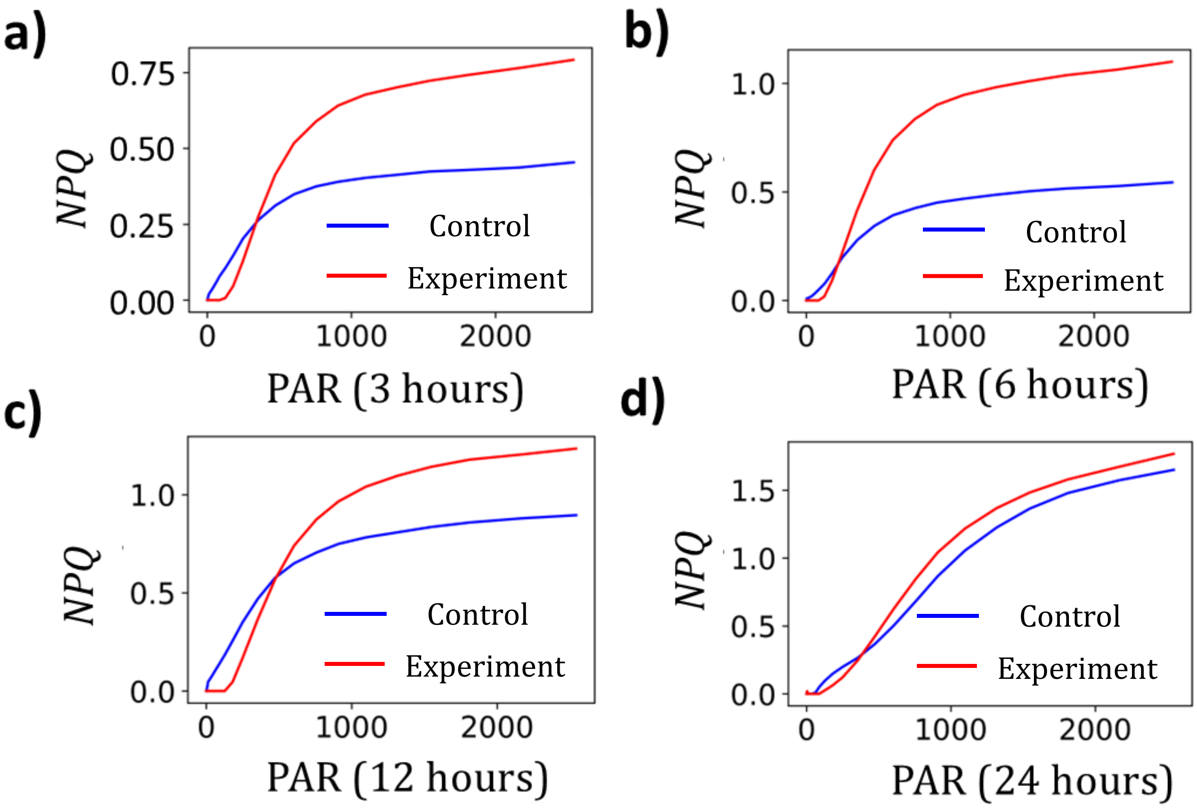


**Fig. S10: *NPQ* distribution for cell samples at different timepoints, under nutrient supplementation. a)** *NPQ* vs PAR distribution for cells after 3 h of initiation in stratification. In all cases, experiment samples represent bioconvecting cells whereas control case represents a suspension where bioconvection is absent. **b, c, d)** *NPQ* vs PAR distribution of samples collected after 6 h, 12 h, and 24 h of mixing via bioconvection (red) and diffusion (blue).

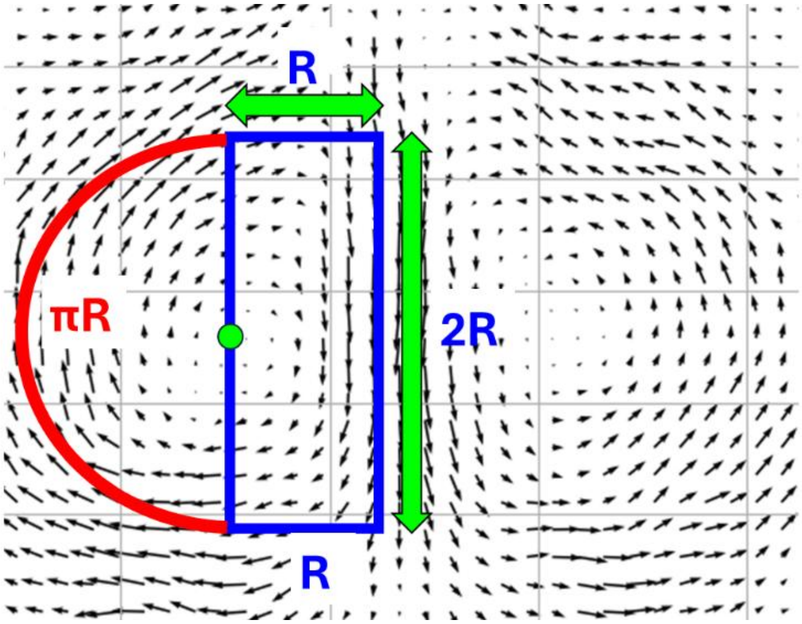


**Fig. S11: Schematic illustration of the geometric path length traversed by gyrotactic cells in bioconvective flows.** This schematic illustrates the approximate geometric path traced by cells during bioconvection, with the trajectory segmented into key transport regions, including the plume center, lateral displacement, and upward migration.
